# Distributed Biomanufacturing of Liquefied Petroleum Gas

**DOI:** 10.1101/640474

**Authors:** Robin Hoeven, John M. X. Hughes, Mohamed Amer, Emilia Z. Wojcik, Shirley Tait, Matthew Faulkner, Ian Sofian Yunus, Samantha J. O. Hardman, Linus O. Johannissen, Guo-Qiang Chen, Michael H. Smith, Patrik R. Jones, Helen S. Toogood, Nigel S. Scrutton

## Abstract

Liquefied Petroleum Gas (LPG) is a major domestic and transport fuel. Its combustion lessens NO_x_, greenhouse gas and particulates emissions compared to other fuels. Propane – the major constituent of LPG – is a clean, high value ‘drop-in’ fuel that can help governments develop integrated fuels and energy policies with low carbon burden, providing solutions to the multi-faceted challenges of future energy supply. We show that bio-LPG (bio-propane and bio-butane) can be produced by microbial conversion of waste volatile fatty acids that can be derived from anaerobic digestion, industrial waste, or CO_2_ via photosynthesis. Bio-LPG production was achieved photo-catalytically, using biomass propagated from bioengineered bacteria including *E. coli, Halomonas* (in non-sterile seawater), and *Synechocystis* (photosynthetic). These fuel generation routes could be implemented rapidly in advanced and developing nations of the world to meet energy needs, global carbon reduction targets and clean air directives.

---

The race to develop economically viable microbial biofuels^1^ is a consequence of a pressing need to reduce carbon emissions, improve air quality and implement renewable and sustainable fuel strategies. Current over reliance on fossil fuels has led to concerns over energy security and climate change. This has driven new policies to restrict greenhouse gas emissions, increase the recycling of waste biomaterials and accelerate the delivery of the bioeconomy.^2, 3^ Effective biofuels strategies would comprise scalable production of transportable and clean burning fuels derived from a robust microbial host, cultivated on renewable waste biomass or industrial waste streams, with minimal downstream processing, and avoiding use of fresh water (**Figure 1**). Embedding production techniques within existing infrastructures for waste processing and fuel distribution would minimise expenditure. Tailoring to specific waste streams would support local economies, waste management, energy self-sufficiency, and carbon reduction in both advanced and developing nations of the world.

**Figure 1.**
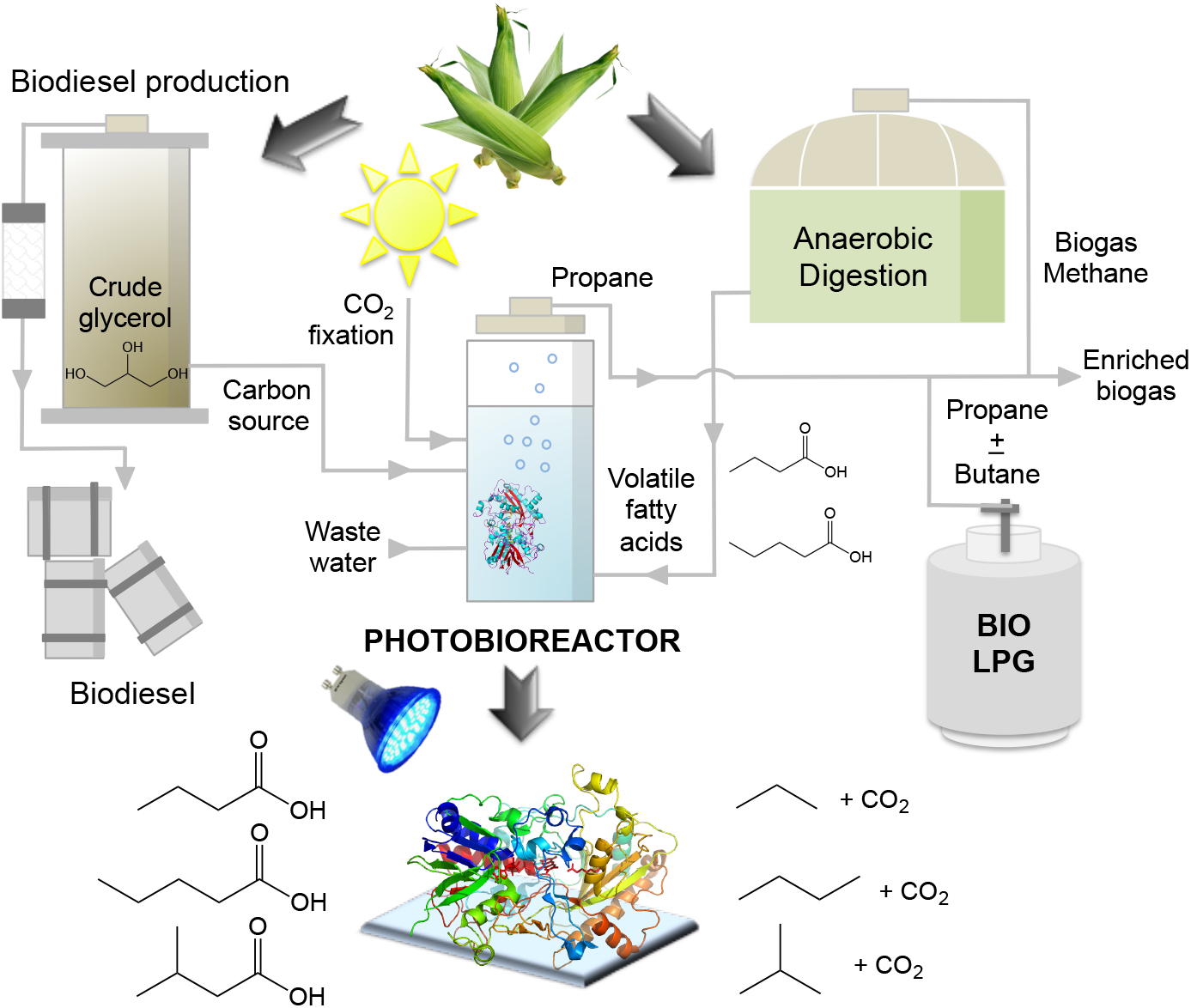
Overview of the bio-LPG strategy. Overview of the bio-LPG production strategy, incorporating a novel photobioreactor design with existing waste feed stock, transportation and distribution infrastructures. The gaseous hydrocarbon blend produced (bio-LPG) is dependent on the relative concentrations of corresponding volatile fatty acids in the feedstock.

Propane is an ideal biofuel. This simple hydrocarbon gas is a highly efficient and clean-burning fuel.^4^ It is currently obtained from natural gas and petroleum refining. Propane is the third most widely used transportation fuel (20 million tons per annum globally) and it is also used for domestic heating and cooking, non-greenhouse gas refrigerants and aerosol propellants. Its ‘drop-in’ nature boosts the calorific value of current methane / biogas supplies, with lower energy requirements for liquefaction and storage.^4^ The only existing commercial production method is the Nesté process, an energy intensive, catalytic chemical conversion of biodiesel waste (glycerol) reliant on natural gas derived H_2_.^5^ No natural biosynthetic routes to propane are known. Engineered biological pathways to propane have been developed based on decarbonylation of butyraldehyde incorporating natural or engineered variants of the enzyme aldehyde deformylating oxygenase (ADO).^6–10^ The low turnover number of ADO (~ 3-5 h^-1^), however, limits implementation of these pathways in scaled bio-propane production.^6,7,9^

The low activity of ADO has stimulated searches for alternative biocatalysts. A novel fatty acid photodecarboxylase (FAP) was described recently that catalyzes blue light-dependent decarboxylation of fatty acids to *n*-alkanes or *n*-alkenes (**Figure 1**).^11, 12^ FAP has a reported reaction quantum yield of greater than 80% and specificity for long chain fatty acids (C14-C18).^11–13^ ADO catalyzes decarbonylation of butyraldehyde (C4) and this can be improved by enzyme engineering.^6^ Likewise, we surmised that FAP could be engineered to decarboxylate the C4 compound butyric acid and other short chain volatile acids, to form propane and other hydrocarbon gases to enable their production at scale.

Here we describe proof-of-concept bio-LPG production techniques that use engineered variants of FAP. These new technologies could be scaled locally for bio-LPG production (e.g. in rural and / or arid communities), and engineered to involve CO_2_ capture, providing alternatives to petrochemical refinery-sourced LPG supplies.

## Light-activated Biocatalysts for Bio-LPG Production

The most suitable FAP enzyme for hydrocarbon gas production was identified by biotransformation assay using cell-free extracts of eight mature N-terminal His_6_-tagged cyanobacterial FAP homologues expressed in *E. coli* (**Table S1**).^11^ With 10 mM butyrate substrate, the highest levels of propane were detected under blue light (455 nm) with the *Chlorella variabilis* NC64A homologue^11–13^ (CvFAP_WT_; 1.65 + 0.61 mg/L lysate). With 10 mM valeric and isovaleric acids, both butane (1.31 mg/L) and isobutane (0.07 mg/L) were generated as detected previously^12^ and low levels of propane were also produced, attributed to the presence of endogenous butyric acid in cell-free extracts. These nascent activities indicate that production of bio-LPG blends, in principle, is feasible using this enzyme. Tunable propane / butane ratios could be achieved by adjusting relative butyric / valeric acid levels. The low gas production levels of CvFAP_WT_ using these volatile short chain carboxylic acid substrates is however a major limitation.

## Enzyme Engineering

Substrate access channels in both CvFAP_WT_^11^ and ADO^14^ are narrow. These adopt a curved architecture to accommodate the long aliphatic chains of C16 / C18 fatty acids and C16 / C18 aldehydes, for CvFAP_WT_ and ADO, respectively. In all other respects, the two enzymes are not related structurally. Of note are residues G462-T484 in CvFAP_WT_ that form part of this substrate access channel (**Figure 2, inset**). A strategy to increase the binding of butyrate is to decrease the competition for the active site by introducing a steric block that impairs the binding of fatty acids of chain lengths greater than C4 / C5. We made a collection of 28 CvFAP variants, targeting residues G462, G455, Y466, V453, T484 and A457 for substitution (**Figure 2**). The side chains of these residues are in close proximity to bound palmitate in the co-crystal structure of CvPAS_WT_ and variants were designed to disfavor long chain fatty acid binding (**Figure 2 inset**).^11^

**Figure 2.**
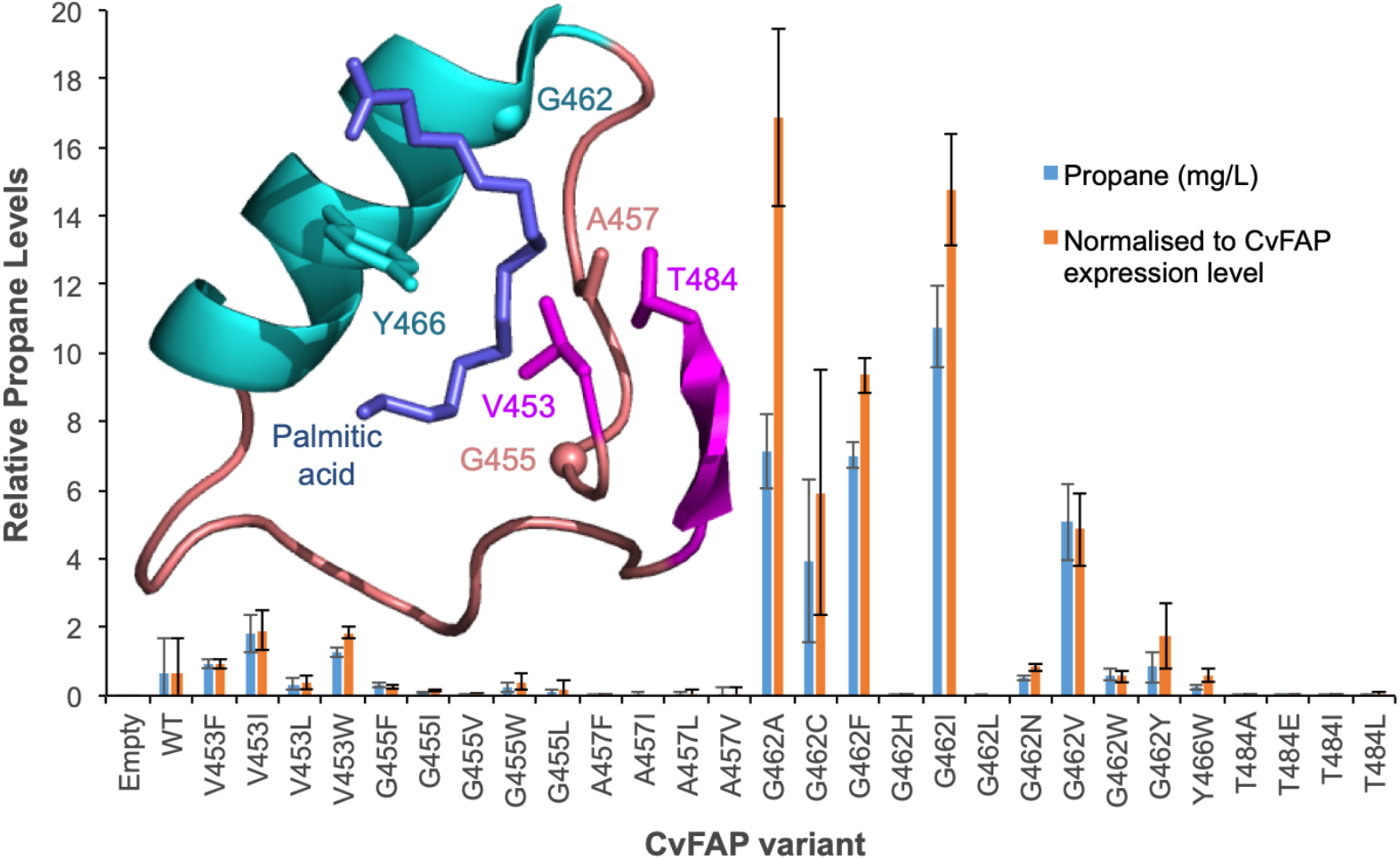
Enzyme engineering for bio-propane production. Comparative propane production of variants of CvFAP in *E. coli.* Transformed *E. coli* cultures were grown at 37 °C in LB medium containing kanamycin (30 μg/mL) to a density of OD_600_ ~ 0.6-0.8. CvFAP gene expression was induced with IPTG (0.1 mM) and cultures were supplemented with 10 mM butyric acid. Triplicate aliquots (1 mL) of cultures were sealed into 4 mL glass vials and incubated at 30 °C for 16-18 h at 200 rpm, illuminated with a blue LED panel. Headspace gas was analysed for hydrocarbon content using a Micro GC. Reactions were performed biological triplicates. Data was normalized by dividing the propane titres (mg/L lysate) by the relative protein concentration compared to the wild type (WT) enzyme (**Figures S1**). Inset: Structure of the palmitic acid binding region of CvFAP (PDB: 5NCC) shown as a cartoon with secondary structure colouring. Palmitic acid and residues targeted for mutagenesis are shown as secondary structure coloured sticks. Residue numbering is taken from the crystal structure (PDB: 5NCC), which is relative to the full deposited amino acid sequence, including the signal sequence. The ‘mature’ sequences in this study are the N-terminally-truncated forms minus the predicted signal sequence (60 amino acids for CvFAP), so residue G462 is actually G402 in the mature sequence.

G462 was substituted for 10 other residues (Val, Asn, Trp, Leu, Cys, Ile, Phe, Ala, His and Tyr). Enzyme synthesis was induced in *E. coli* cultures exposed to blue light (**Figures S1-S2**), and propane production was measured and normalized according to the relative expression level of each variant (SDS PAGE; **Figure S3**). Variant CvFAP_G462V_ showed a 7-fold increase in propane titre (5.07 + 1.12 mg propane per L live culture per day; **Figure 2; Table S2**) compared to CvFAP_WT_.

With (iso)valeric acids (C5), CvFAP_G462V_ (iso)butane production was 2-fold greater than for CvFAP_WT_ (2.52 and 1.31 mg/L culture, respectively). Further increases in propane production were achieved with variants G462I, G462F and G462A (1.9-3.5 fold greater than G462V; **Table S2**). Increased propane production correlates with binding constants determined from molecular docking simulations (Autodock Vina^15^) for CvFAP_WT_ and variants G462V and G462I, in which a 30 – 50-fold decrease in binding constants (relative -ΔG_binding_ 1.4–1.5-fold; **Table S3**) with palmitate, and a small increase with butyrate binding (1.4-fold; **Figure 3**) were predicted. Overall, mutagenesis of residue G462 leads to as much as a 25-fold increase in propane titre from butyric acid. With the exception of G455I, variants at other positions (G455, Y466, V453, T484, G455 or A457) produced less propane than CvFAPWT (**Figure 2**). Thus, a range of biocatalysts has been produced with side chain replacements at residue 462 being more effective at enhancing short chain fatty acid decarboxylation than at other targeted positions.

**Figure 3.**
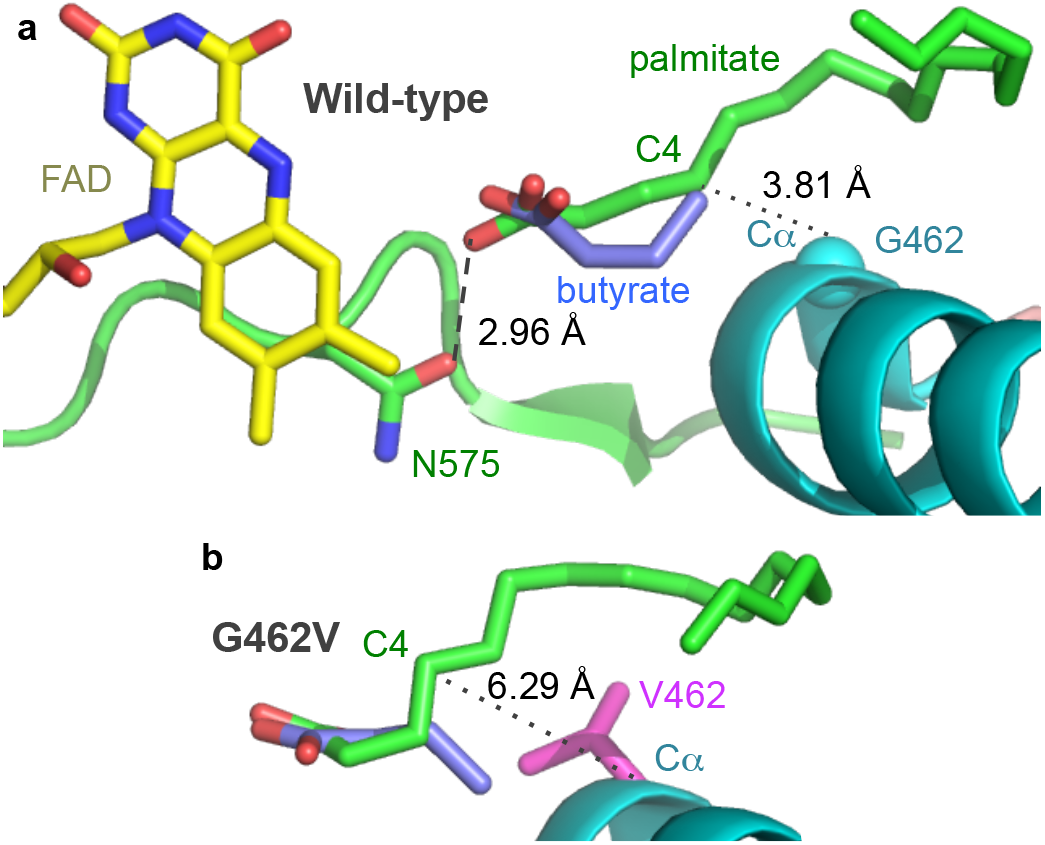
Active site of cvFAP. Models of butyrate and palmitate in the active site of a) wild-type and b) G462V variant of CvFAP. The position of palmitate in the wild-type enzyme is crystallographically determined (PDB:5NCC). The positions of the remaining ligands were determined by Autodock Vina,^17^ and mutagenesis to G462V was simulated using SwissPDBViewer 4.10.^46^ In principle, increasing the side chain size at position 462 should decrease the volume of the substrate access tunnel and disfavour energetically the binding of palmitate. These models suggest the distance between the Cα-atom of G462V and the C4 atom of the substrate is significantly increased due to the presence of the isopropyl group of valine in the place of hydrogen in glycine. This repositioning of palmitate to a less favourable orientation relative to the wild-type complex could explain the decrease in the predicted *K*_b_ for the variants. The exception is variant G455I that showed a near 2-fold increase in palmitate *K*_b_. In both panels, the protein is shown as a cartoon with secondary structure coloring, with selected residues shown as sticks. FAD, palmitate and butyrate are shown as atom-colored sticks with yellow, green and blue carbons, respectively. The dashed line shows a hydrogen bond between palmitate and the wild-type enzyme, while the dotted lines only highlight the modeled ligand-G/V462 residue distances.

A possible limitation to propane production in live cells is cell permeability to the substrate (butyric or other volatile acid). In live cell cultures, the highest propane levels were detected with 10 mM butyric acid (7.53 + 0.29 mg/L culture). Higher concentrations affected the culture pH with concomitant cytotoxic effects (**Figure S4**). A potential solution is to increase butyrate uptake by the addition of cell permeabilization agents (e.g. Triton X-100 and / or sucrose)^16^ but this led to only a modest increase in propane titres (1.6-fold; **Table S4**). Another approach is to stimulate the butyrate uptake transporter atoE (part of the *atoDAEB* small chain fatty acid catabolism operon) with acetoacetate.^17, 18^ Supplementation with ethyl- or methyl acetoacetate produced *ca* 2- and 2.8-fold more propane, respectively. Co-expressing a recombinant atoE transporter with CvFAPG462V did not increase propane titres to any major extent (1.3-fold; **Table S4**), even in the presence of acetoacetate.^17^ Butyrate uptake and toxicity will be specific to the host microorganism and for distributed biomanufacture is best optimized with the selected industrial strain (see below).

We observed varying effects on propane production according to the plasmid backbone used (pETM11 or pET21b) and the size and position of the associated poly(His)-tag (whether N- or C-terminal). Both plasmids contain the same ColE1 origin of replication and a T7lac promoter. However, pETM11 incorporates a TEV protease-cleavable N-His_6_-tag, whereas pET21b incorporates a shorter C-terminal His_6_-tag. We observed a 6.4-fold increase in propane production using CvFAP_G462V_ contained in plasmid pET21b compared to pETM11 (48.31 + 2.66 mg/L culture). This was increased further (97.1 + 10.3 mg/L; **Table S5**) by adding ethyl acetoacetate. This highlights the need to explore multiple plasmid backbones and the location / size of protein tags for gas production *in vivo*. Since pET21b supported the highest propane yields, this plasmid backbone was used in subsequent studies (below).

## Tunable Bio-LPG Blends

Photodecarboxylation of other short chain fatty acids (butyric, isobutyric, valeric, 2-methylbutyric and isovaleric acid) was investigated with wild-type and four CvFAP variant enzymes (G462V/A/I/F). Propane, butane and isobutane was produced (**Figure 4a**). Gas levels were greater with variant G462I (compared to G462V), particularly with the branched chain substrates (isovaleric and 2-methylbutyric acids; 5-8-fold higher; **Table S6**). With CvFAP_G462_, propane and butane production from linear substrates (butyric and valeric acid) were less than 2-fold higher than CvFAP_G462V_. Variants G462V and G462A generated similar levels of propane and butane, but with a greater variation in hydrocarbon titre seen with the G462A variant (**Figure 4a; Table S6**). The G462 position is clearly important in conferring activity with other short chain carboxylic acids (other than propane) required to make LPG-blends. As variant G462V generated similar titres of propane and butane it was taken forward as a suitable biocatalyst to develop tunable bio-LPG blends.

**Figure 4.**
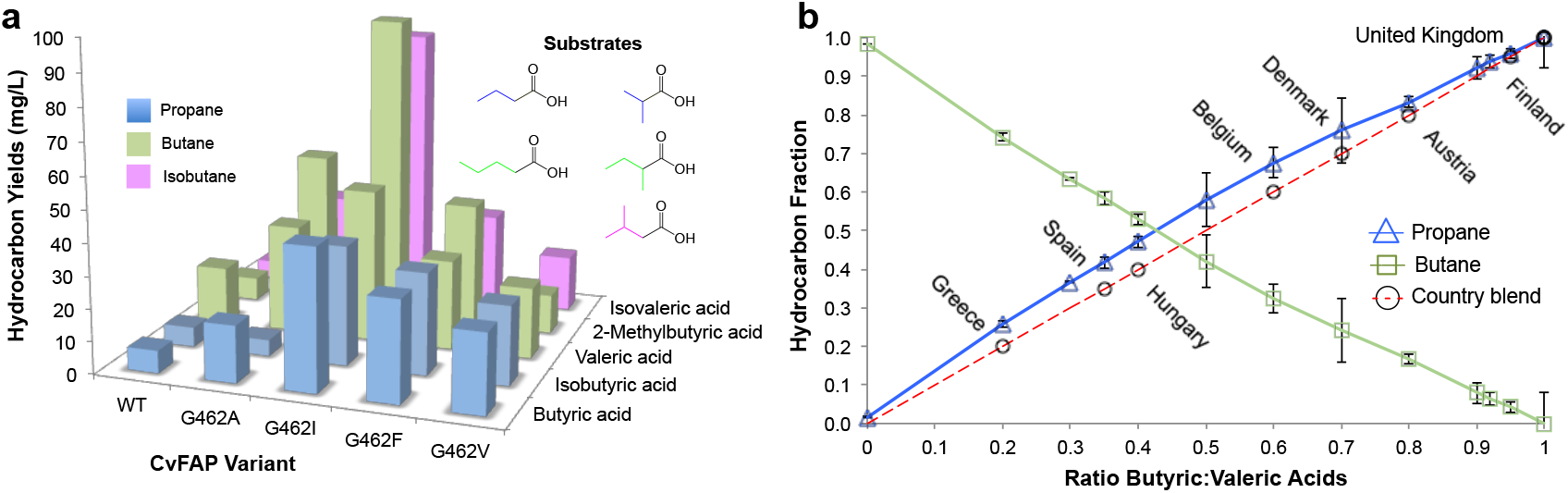
Tunable bio-LPG blends. *In vivo* gaseous hydrocarbon production by wild type and variant CvFAP in *E. coli* st. BL21(DE3)ΔyqhD/ΔyjgB. Effect of a) CvFAP-pETM11 variant^a^ and b) butyric:valeric acid blends with CvFAP_G462V_-pBbA1c on hydrocarbon production. Cultures (20 mL) were grown in LB medium containing kanamycin (50 μg/mL) at 37 °C until OD_600_ ~ 0.6-0.8. Recombinant protein expression was induced with IPTG (0.1 mM) followed by culture supplementation with fatty acid substrates (10 mM) after 1 h at 30 °C. Triplicate aliquots (1 mL) of cultures were sealed into 4 mL glass vials and incubated at 30 °C for 16-18 h at 200 rpm, illuminated with a blue LED panel. Headspace gas was analysed for hydrocarbon content using a Micro GC. ^a^All reactions designed to generate butane and isobutane also produced ~ 2% propane. The numerical data for both panels can be found in **Tables S6-S7**.

The most common gases found in LPG blends are propane and n-butane. Blends may also contain isobutane, ethane, ethylene, propylene, butylene and isobutylene. The exact composition of LPG gases is country-specific, and can vary between seasons.^19^ In the UK, LPG is 100% propane, while in Italy the propane:butane ratio varies from 90:10 to 20:80 (**Figure 4b**). As CvFAP_G462V_ can generate both propane and butane at similar titres, we investigated the possibility of production of country-specific bio-LPG blends *in vivo* by varying the ratio of externally supplied butyric:valeric acids. There was a remarkably close correlation between the proportions of butyric:valeric acid in feedstock and the respective propane:butane concentration in the culture headspace (**Figure 4b; Table S7**). The relative ease at which highly tunable bio-LPG blends were generated shows the potential applicability of this process whereby country-specific requirements can be met by a simple manipulation of the volatile fatty acid feed ratio.

## Distributed Manufacture of Bio-LPG from Waste Biomass Streams

Microorganism-derived fuels and chemicals production is costly and places high demands on both capital and operational expenditures. Typically, steel-based bioreactors with complex monitoring systems are used. These are associated with high running costs (e.g. energy-intensive aeration, mixing and downstream processing). Sterilisation of equipment is needed to minimise microbial contamination and growth under aseptic conditions is used. There are also environmental concerns over waste processing and disposal, and production methods use large quantities of clean water. These multi-faceted issues can drive up production costs. We tackled these challenges at the outset by selecting *Halomonas* as the production host as it is proven to grow under non-sterile conditions with minimal risk of contamination.^20^

*Halomonas* is both halophilic and alkaliphilic and can grow in NaCl concentrations as high as 20% (w/v) and pH as high as 12. Continuous cultures have been grown for over three years in industrial-scale vessels for the biomanufacture of polyhydroxyalkanoates. This has been achieved at >1,000 tonnes scale, with no decline in growth potentials.^21^ Seawater and recycled water can be used without sterilisation, conserving fresh water and reducing energy expenditure. Major capital savings and efficiency gains can therefore be made as bioreactors can be constructed using low cost materials (e.g. plastics, ceramics and cement). Scaled production of polyhydroxyalkanoates using *Halomonas* is at *ca* 65% cost saving compared to *E. coli*.^22, 23^ This suggests that distributed biomanufacture of Bio-LPG in *Halomonas* could similarly benefit from major capital and operational expenditure savings.

A Halomonas-compatible plasmid construct pHal2 containing CvFAP_G462V_ was generated using a broad host range pSEVA plasmid,^24^ containing an IPTG-inducible T7-like promoters (MmP1-*lac*O-RiboJ-SD; **Figure 5a**)^25^ (**Figure S5**). Gas production studies were performed in a non-sterile high salt minimal medium (6% NaCl) using *Halomonas* strain XV12, which contains a chromosomal copy of the equivalent T7-like RNA polymerase.^26^ *Halomonas* live cells expressing pHal2-CvFAPG462V generated propane at levels similar to that seen with *E. coli* constructs at 10 mM butyrate (78.9 + 14.1 mg/L culture; **Figure 5b**), which was 8-fold higher than the equivalent wild-type construct. The switch in host from *E. coli* to *Halomonas* therefore has not significantly diminished propane titres, even though the *fap* gene has not been optimised for codon usage in *Halomonas*. Studies using cell permeabilization or atoB transporter stimulation reagents showed no significant effects on propane titres (**Table S8**), in contrast to equivalent reactions with *E. coli.* This may be related to cell wall and phospholipid adaptations for growth under halophilic conditions.^27^ The lack of stimulation by ethyl acetoacetate was unexpected given that putative atoE genes are present in the *Halomonas* genome.

**Figure 5.**
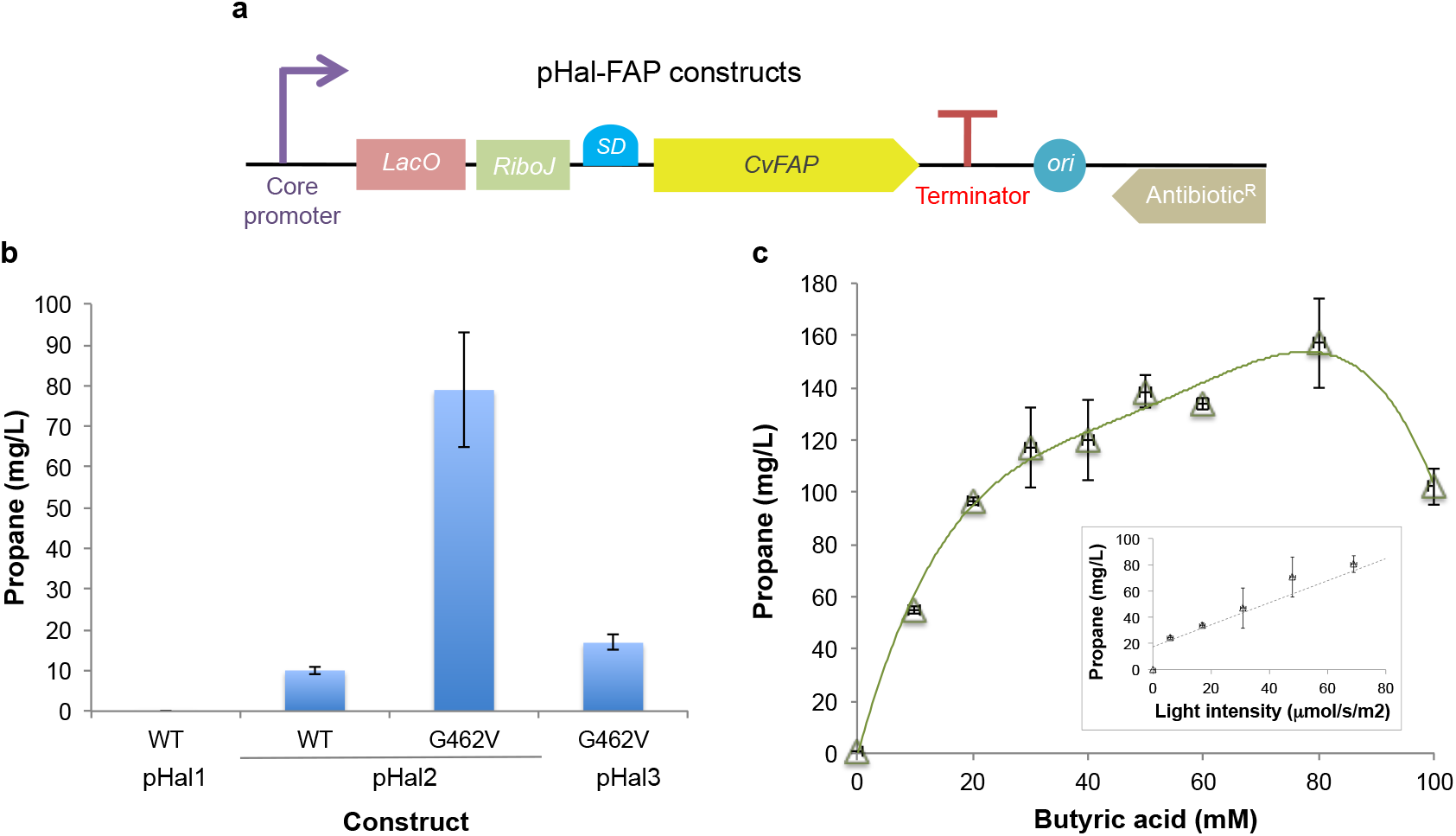
Propane production *by Halomonas* strain XV12. a) Schematic and b) propane production of *Halomonas* constructs expressing CvFAP. c) Effect of butyric acid concentration on propane production by pHal2_G462V_. Inset: effect of light ‘intensity’ (i.e. photosynthetic photon flux density (PPFD) up to 140 μE) on propane production by the same construct. Constructs (all non-His_6_-tagged) were generated in the following modified plasmids: pSEVA321 (pHal1-FAP_WT_ and pHal3-FAPo462v and pSEVA441 (pHal2-FAP_WT_ and pHal2-FAP_G462V_). Cultures were grown in phosphate buffered YTN6 medium spectinomycin (pHal2-FAPG462V; 50 μg/mL) or chloramphenicol (pHal1- and pHal3 constructs; 34 μg/mL) for 5 h at 37 °C and 180 rpm. Recombinant protein expression was induced with IPTG (0.1 mM) at an OD_600_ ~ 1.6), and cultures were supplemented with butyric acid (0-100 mM). Triplicate aliquots (1 mL) of cultures were sealed into 4 mL glass vials and incubated at 30 °C for 16-18 h at 200 rpm, illuminated with a blue LED panel. Headspace gas was analysed for hydrocarbon content using a Micro GC.

*Halomonas* cultures displayed a relatively high tolerance to butyric acid (80 mM) in the presence of buffering salts. Propane titres reached 157.1 + 17.1 mg/L culture under these conditions (**Figure 5c**); the highest reported production to date. These titres are *ca* 9x greater than titres reported for *E. coli* containing engineered pathways that contain ADO, and 5x greater than *E. coli* containing CvFAP_G462V_ and fed with butyrate.^6, 7, 9^ In *Halomonas*, access of light to the biocatalyst was found to be a limiting factor as revealed by the linear dependence of propane titre on light intensity (**Figure 5c inset**). Optimal light intensity is a balance between competing factors. These include high cell densities to increase photobiocatalyst supply, poor penetration of light at high cell densities, and adverse effects on cell viability of high light intensities at high cell densities. Light intensity however can be controlled in scaled photobioreactor units through appropriate design.

Under laboratory conditions, the switch to *Halomonas* improved propane titres substantially. For production in the field, engineered strains would need to use waste biomass feedstock streams and seawater / recycled water at scale. This requires aerobic growth on simple carbon sources (e.g. sugars, glycerol) in high salt concentrations with minerals, vitamins and butyric (or other small organic) acids. Seawater is a cost-effective natural source of mineral and salt (3.5% w/v salinity), while clarified wastewater streams provide an abundant alternative for inland sites. Further sea salt supplementation to the required salinity and mineral content at high alkalinity will sanitise the medium bypassing any requirement for sterilisation, as almost all competing microorganisms are unable to grow under these conditions. Vitamins can be supplied, for example, from autolysed spent brewery yeast, an abundant waste product, or similar. For bacterial growth, raw biodiesel waste is a cost-effective carbon source^28^ as a low value product composed primarily of glycerol (60-70%), salts, methanol and residual vegetable oils (**Figure 1**).^29^ Butyric acid and other volatile acids can be sourced from anaerobic digestion (AD) of lignocellulosic agricultural biomass and food waste.

The potential to scale Bio-LPG production using waste feed stocks was investigated at laboratory scale (400 mL) using a cyanobacterial flatbed photobioreactor. Comparative non-sterile aerobic fermentations were performed between ‘clean’ (laboratory grade reagents) and ‘crude’ (filtered seawater and biodiesel waste glycerin) media components in batch culture with online headspace monitoring for propane production. The presence of seawater and biodiesel waste impurities showed only a minor effect on *Halomonas* growth (**Figure 6a**) and the maximum rate of propane production (82 vs 100 mg/g cells/day; **Figure 6b**). This suggests that inexpensive and abundant waste feed stock streams and seawater can indeed be exploited when considering the design of large-scale bioreactors for renewable bio-propane production.

**Figure 6.**
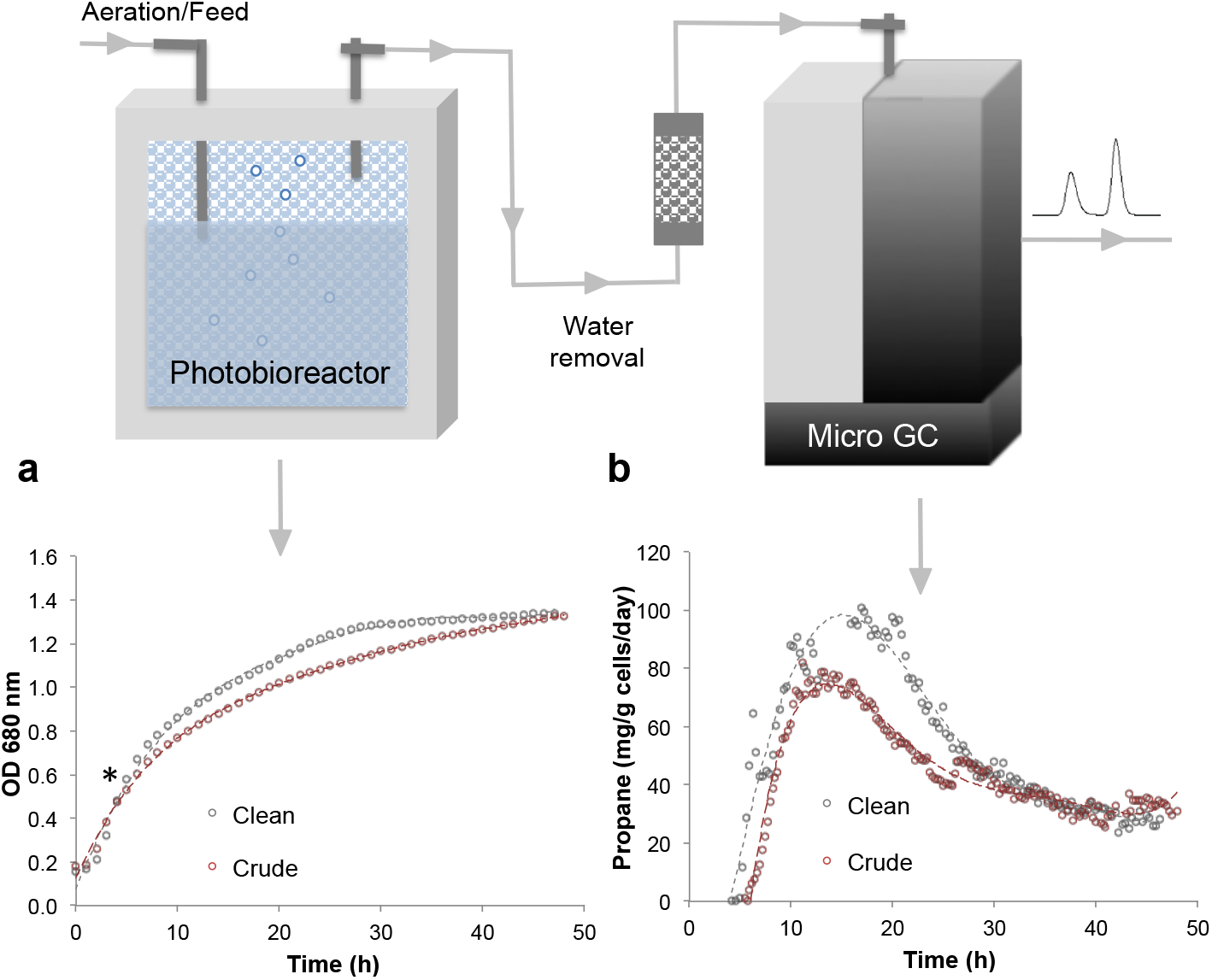
Cultivation of *Halomonas* expressing pHal2-FAP_G462V_. The figure shows a) culture growth (OD 680 nm) and b) propane production. Cultures (400 mL) were grown in high salt glycerol medium at pH 6.8 containing 50 μg/mL spectinomycin and 0.5 mL/L antifoam. ‘Clean’ fermentations included purified glycerol/NaCl, while ‘Crude’ ones contained biodiesel waste glycerin with sea water supplemented with additional NaCl (6% total salinity). Conditions were maintained at 30 °C with 65-100% stirring, an airflow rate of 1.21 L/min, automated pH maintenance, culture optical density monitoring and ambient room lighting until mid-log phase (4-5 hours). Recombinant protein expression was induced with IPTG (0.1 mM), followed by the addition of sodium butyrate (60 mM pH ~6.8) and blue light exposure (1625 μE), and maintained for ~48 h. Propane production was monitored at 15-20 minute intervals by automated headspace sampling using a Micro GC.

Online monitoring of the headspace during *Halomonas* cultivation revealed a peak and then a steady decline in propane production over time (**Figure S6**). This is commonly seen with plasmid-borne constructs (**Figure 6b**), and is attributed to plasmid instability and / or loss. Genomic integration of the G462V variant (or other variants) for constitutive expression would safeguard against this loss for prolonged scaled production. Scaled photobioreactors could be coupled with existing headspace extraction / liquefaction technologies and distribution infrastructures for LPG. Implementation of such a strategy at scale could transform distributed manufacture of gaseous hydrocarbon fuels from waste biomass and wastewater / seawater.

## Bio-propane Production by Photosynthetic Carbon Capture

An ideal energy strategy would be to develop a sustainable carbon neutral fuel, whereby its combustion emissions (CO_2_) could be recycled as the carbon source for the production of further biofuel. Existing technologies of carbon capture and storage (CCS) have been implemented to significantly reduce emissions, such as at fossil fuel electricity generating plants, cement, steel and chemical works. The International Energy Agency has estimated that CCS could potentially contribute to a 19% reduction in CO_2_ emissions by 2050.^30^

A natural (microbial) carbon capture solution would take advantage of the photosynthetic ability of microorganisms such as cyanobacteria to fix CO_2_ into organic carbon.^31^ The cyanobacterium *Synechcocystis* PCC 6803 is an ideal host because it grows rapidly and it is genetically tractable,^32, 33^ tolerant to abiotic stress,^34^ and conditions for optimizing its growth are well understood.^35, 36^ The photobiological conversion of CO_2_ into medium chain-length fatty acids^37^ and long chain hydrocarbons^38^ in *Synechcocystis* PCC 6803 has been described, the latter utilising an ADO- or FAP-based system. Thioesterase A from *E. coli* was incorporated, which catalyses the conversion of fatty acyl-ACP to free fatty acids, in addition to knocking out the native fatty acyl ACP synthase gene (Δaas) to minimise the reverse reaction (**Figure 7a**).^38^ Together these changes increased the availability *in vivo* of free fatty acid precursors,^38^ enabling hydrocarbon biosynthesis to occur direct from CO_2_, instead of via external carbon source addition.^38^

**Figure 7.**
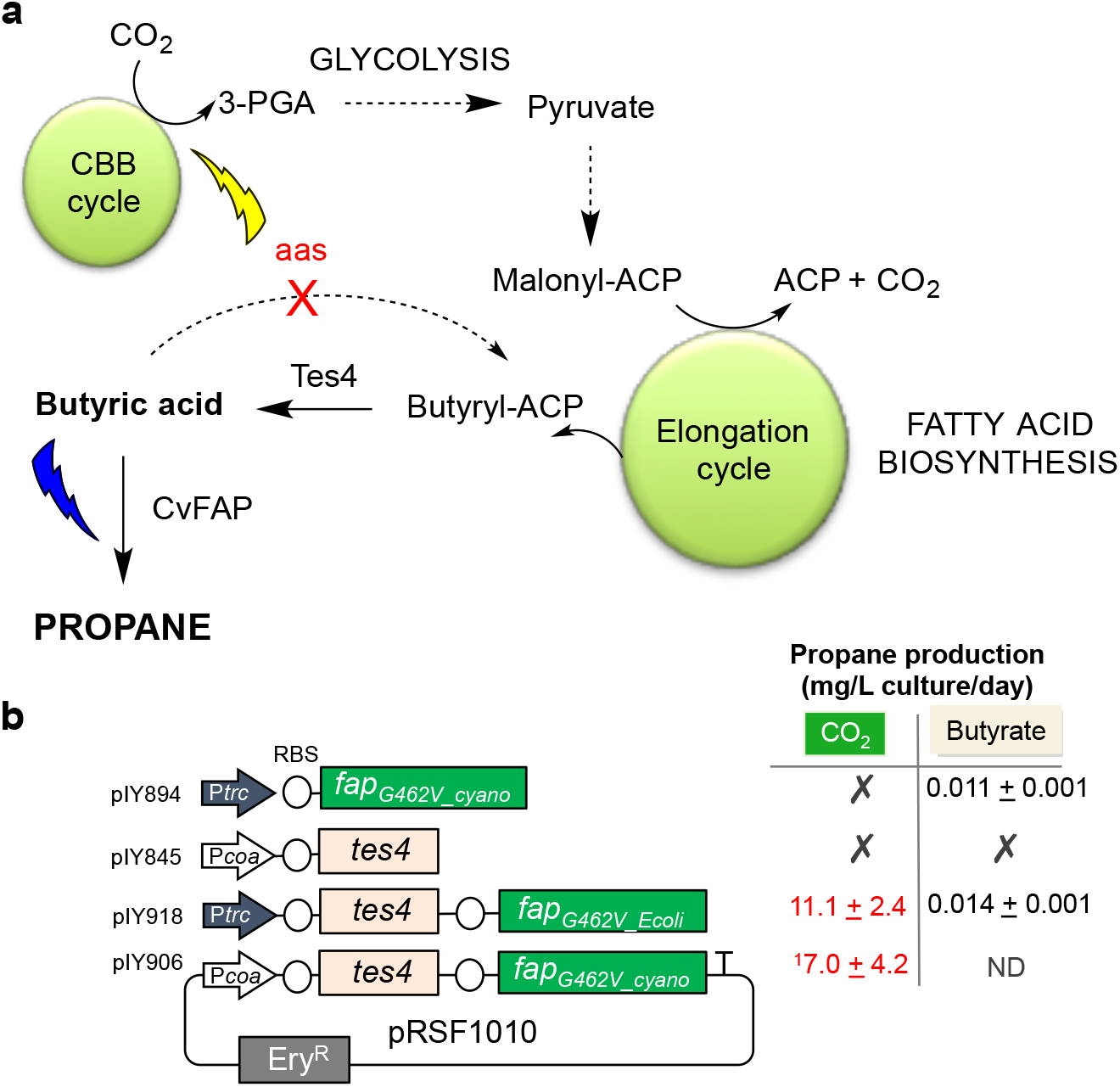
Photoautotrophic production of propane. Metabolic engineering enabling the conversion of CO_2_ to propane via the up-regulation of butyric acid production in vivo, a) Engineering scheme and b) propane production from plasmid constructs in *Synechocystis* Δ*aas* from CO_2_ or butyrate feeding. Culture growth was performed in batch mode (sealed vials) in the presence or absence of 10 mM butyrate, and under standard photosynthetic conditions in a photobioreactor (see Methods). The following constructs were generated: *i*) pIY894: Ptrc::*fap_G462V_cyano_*; *ii*) pIY918: Ptrc::*tes4, fap_G462V_Ecoli_*; *iii*) pIY906: Pcoa::*tes4, fap_G462V_cyano_*; and *iv*) pIY845: Pcoa::*tes4*. Ptrc = *E. coli* promoter lacking the *lacI*, making it a constitutive promoter. Red numbers indicate propane production in a photobioreactor, while black numbers were obtained from batch cultures in the presence of 10 mM butyrate. ^1^Propane production detected in non-induced cultures. Each data point is an average of biological triplicates. CBB cycle = Calvin–Benson–Bassham cycle; 3-PGA = 3-phosphoglycerate; ACP = acyl carrier protein; aas = acyl-acyl carrier protein synthase; Tes4 = acyl-ACP thioesterase; Ery^R^ = erythromycin resistance gene. ND = not determined as because the higher expression of the cyanobacterial codon optimised gene CvFAP_G402V_cyano_ appeared to be toxic and reduced cell growth rates.

We incorporated an *E. coli* or *Synechocystis* codon-optimised gene for CvFAPG462V into *Synechocystis Δaas* strain and / or *tes4* (TesA homologue from *Haemophilus influenza)* under the control of a cobalt (II)-inducible (Pcoa)^39^ or constitutive promoter (Ptrc/lacI^-^; **Figure 7b**). Cultures were grown under standard photosynthetic conditions to moderate cell density (OD 720 nm = 1) before aliquots were sealed in glass vials for 24-48 h under blue light with or without exogenously added butyric acid. As expected, *Synechocystis* wild-type and *Δaas* +/- Tes4 control strains did not produce any detectable propane, and Tes4-containing constructs showed a near 3-fold increase in butyrate titres (**Figure 7b**). Low levels of propane production were only detected in CvFAP_G462V_-containing batch cultures externally supplied with butyrate (pIY894:Ptrc-CvFAP_G462V_cyano_ and pIY918: Ptrc-tes4-CvFAP_G462V_Ecoli_; **Figure 7b**).

*Synechocystis* Δ*aas* strain pIY918 was cultivated in the photobioreactor under more standard photosynthetic conditions (aeration, CO_2_ supply, pH control) with supplementary blue light exposure, to see if propane production occurs from CO_2_-derived butyric acid. This constitutively expressed culture showed moderate propane production (11.1 + 2.4 mg propane/L culture/day), which is equivalent to ~12.2 + 2.6 mg propane/g cells/day (**Figures 7b and S7**). This is the first demonstration of fully autotrophic propane production from CO_2_. Propane titres are around 8-fold lower than the equivalent *Halomonas* pHal2-FAPG462V host. Proof-of-concept for autotrophic propane production from CO_2_ has now been established and this will enable optimisation in further work. Autotrophic growth with butyrate feeding is also possible, which is considerably less expensive than traditional fermentation media. Moreover, photosynthetic production of butyrate from industrial CO_2_ by the Tes4-containing *Δaas* knockout strain could provide an alternative to butyrate supply for the *Halomonas* pHal2-FAP_G462V_ chassis. This would enable CO_2_ (photosynthetic carbon capture) as well as waste biomass (e.g. anaerobic digestion) to be used as a source of butyrate for the *Halomonas* propane production strain.

## Conclusions

We have described platform technologies for the production of propane and other bio-LPG components based on the exploitation of light-activated biocatalysts and laboratory / industrial microbial strains. These technologies will enable distributed biomanufacture of gaseous hydrocarbons at low cost using waste industrial / agricultural feedstocks, including CO_2_. The use of *Halomonas* will lead to reduced biomanufacturing costs for scaled gas production. Bio-LPG production coupled to photosynthetic carbon capture enhances further these biomanufacturing capabilities. A blending of these approaches (e.g. the engineering of halophilic cyanobacteria, such as *Euhalothece^40^)* would unite the benefits of low-cost production with carbon capture. Scaled biomanufacture using these technologies offers solutions to urgent global energy needs, waste management, reduction of CO_2_ and the conservation of fresh water.

## Methods

### Materials and equipment

All chemicals and solvents were purchased from commercial suppliers and were of analytical grade or better. Media components were obtained from Formedium (Norfolk, UK). Gene sequencing and oligonucleotide synthesis were performed by Eurofins MWG (Ebersberg, Germany). All oligonucleotide sequences can be found in the Supplementary **Tables S9-S11**. The mounted high-power blue LEDs and LED drivers were from Thorlabs (Ely, U.K.), with spectra centered at 455 nm (bandwidth (FWHM) 18 nm,1020 mW typical output) and 470 nm (FWHM 25nm, 710 mW typical output). The domestically sourced white light LED (Integral) had a 25W power (2060 lumens). The photobioreactor was a thermostatic flat panel FMT 150 (400 mL; Photon Systems Instruments, Czech Republic) with integral culture monitoring (OD 680 nm), pH and feeding control and an LED blue light panel (465 nm; maximum PPFD = 1648 μE photons).

*E. coli* strain BL21(DE3) was modified by chromosomal deletion of two aldehyde reductase genes *yqhD* and *yjgB* (BL21(DE3)ΔyqhD/ΔyjgB/Kan^R^; GenBank: ACT44688.1 and AAA97166.1, respectively) as described previously.^6^ The kanamycin selection gene was removed using the Flp-mediated excision methodology (BL21(DE3)ΔyqhD/ΔyjgB).^41^ *Synechocystis* sp. PCC 6803 was modified by chromosomal deletion of the acyl-ACP synthetase (*aas*) encoding gene as described previously.^37, 38^ *Halomonas* strains TD01^21^ and TQ10, and modified pSEVA plasmids have been described previously.^26^ *Halomonas* strain XV12 is a modified version of the TQ10 strain, which had been cured of a recombinant plasmid.

### Gene synthesis, sub cloning and mutagenesis

Codon-optimised gene synthesis of the following N-terminally truncated (ΔN) FAP enzymes was performed by GeneArt (Thermo Fisher; **Table S12**): CvFAP_WT_ from *Chlorella variabilis* NC64A^11^ (Genbank: A0A248QE08; ΔN-61); CcFAP from *Chondrus crispus* (UniProt: R7Q9C0; ΔN-50 amino acids truncated), ChFAP from *Chrysochromulina* sp. (UniProt: A0A0M0JFC3), CmFAP from *Cyanidioschyzon merolae* (UniProt: M1VK13; ΔN-64), CrFAP from *Chlamydomonas reinhardtii* (UniProt: A8JHB7; ΔN-31), CsFAP from *Coccomyxa subellipsoidea* (UniProt: I0YJ13; ΔN-43), GpFAP from *Gonium pectorale* (UniProt: A0A150GC51; ΔN-38) and PtFAP from *Phaeodactylum tricornutum* (UniProt: B7FSU6)^11^. Each gene was sub cloned into pETM11 (*Nco*I-*Xho*I), incorporating a TEV protease cleavable 78 bp N-His_6_-tag for rapid protein purification. An additional codon-optimised synthesised gene was synthesised by GeneArt, namely the short chain fatty acid transporter atoE from *E. coli* (UniProt: P76460) with its native OXB1 promoter in pET21b without a C-terminal His_6_-tag (www.oxfordgenetics.com; **Table S12**).^17, 18^ The gene encoding thioesterase Tes4 from *Bacteroides fragilis* (UniProt: P0ADA1) was obtained from plasmid pET-TPC4, as described previously.^6^

Variant CvFAP_G462V_ was generated by site-directed mutagenesis of the wild-type construct in pETM11 using the QuikChange whole plasmid synthesis protocol (Stratagene) with CloneAmp HiFi PCR premix (Clontech). The additional variants G462N/W/L/C/I/F/A/H/Y were generated using the Q5 and QuikChange site directed mutagenesis kits, according to the manufacturer’s protocol (New England Biolabs and Novagen). In each case, PCR products were analysed by agarose gel electrophoresis and gel purified using the NucleoSpin Gel and PCR Clean-up kit (Macherey-Nagel). Constructs were transformed into *E. coli* strain NEB5α (New England Biolabs) for plasmid recircularization and production. The presence of the mutations was confirmed by DNA sequencing followed by transformation into *E. coli* strains BL21(DE3) and BL21(DE3)ΔyqhD/ΔyjgB^6^ for functional expression studies.

### Molecular modelling

Substrates palmitic and butyric acid docked into chain A of the crystal structure of the palmitic acid bound CvPAS structure 5NCC using Autodock vina.^15^ AutoDock Tools 1.5.6 was used to assign non-polar hydrogens and prepare input files. A cubic search volume with sides of 15 Å was defined with the coordinates of C6 of palmitic acid as the centre, and an exhaustiveness of 50 was used to generate 20 conformations, out of which the lowest-energy conformation with the substrate in the correct orientation (carboxylate pointing towards the FAD) was selected. Mutations were performed in SwissPDBViewer 4.10,^42^ using the exhaustive search function to identify the best rotamer for the mutated residue.

### *Escherichia coli* multi-enzyme construct generation

Dual gene construct CvFAP_G462V_-atoE was generated by ligation of PCR amplified CvFAP_G462V_ into the existing atoE-pET21b construct by In-Fusion cloning, with each gene controlled by its own promoter (T7 and OXB1, respectively). Additional constructs of N-His_6_-CvFAP_G462V_ were generated in plasmids pET21b and pBbA1c^43^ by PCR-mediated In-Fusion cloning. Constructs were transformed into *E. coli* strain NEB5α, BL21(DE3) and BL21(DE3)ΔyqhD/ΔyjgB^6^ for functional expression studies.

### *Synechocystis* construct generation

Two versions of the non-His_6_-tagged *C. variabilis* CvFAP_G462V_ gene with identical amino acid sequences were constructed in *Synechocystis* sp. PCC 6803 (*fap_G462V_Ecoli_* and *fap*_*G462V*_cyano_), differing by applying codon optimisation for *E. coli* and *Synechocystis*, respectively. For*fap*_*G462V*_cyano_, plasmid pIY505 (pJET-‘FAP’)^38^ variant G462V was generated by site-directed mutagenesis using the QuikChange whole plasmid synthesis protocol. To construct pJET-*fap_G462V_Ecoli_*, the non-His_6_-tagged gene in pETM11 was amplified by PCR and cloned into the blunt-ended pJET1.2 plasmid. The gene encoding Tes4 was amplified from construct pET-TPC4^6^ and ligated into blunt pJET1.2 plasmid. To clone the mutated *fap* genes and/or *tes4* genes into the erythromycin resistant RSF1010 plasmid, the Biopart Assembly Standard for Idempotent Cloning (BASIC) method was used as described previously.^37, 38, 44^ Gene expression was controlled using either the cobalt-inducible Pcoa or constitutive Ptrc (no lacI) promoters. Prefix and suffix linkers used to create the plasmids are listed in Tables S11-S12. The following constructs were generated: *i*) pIY894: Ptrc-*fap*_*G462V_cyano*_; *ii*) pIY918: Ptrc-*tes4, fap_G462V_Ecoli_*; *iii*) pIY906: Pcoa-*tes4, fap_G462V_cyano_*; and *iv*) pIY845: Pcoa-*tes4*. Plasmid assembly was validated by DNA sequencing.

Plasmids were transformed into the *E. coli* helper/cargo strain (100 μL; *E. coli* HB101 strain carrying the pRL623 and RSF1010 plasmids), conjugal strain (*E. coli* ED8654 strain carrying pRL443 plasmid)^45^ and *Synechocystis* sp. PCC 6803 lacking acyl-ACP synthetase (encoded by *slr1609*; Δaas strain; OD_730_ ~1) using the tri-parental conjugation method described previously.^37, 38^ Each strain had been pre-treated by washing with LB and BG11-Co medium for *E. coli* and *Synechocystis*, respectively, to remove antibiotics. The mixture was incubated for 2 h (30 °C, 60 μmol photons m^-2^ s^-1^), then spread onto BG11 agar plates without antibiotic, and incubated for 2 d (30 °C, 60 μmol photons m^-2^ s^-1^). Cells were scraped from the agar plate, resuspended in 500 μL of BG11-Co medium, and transferred onto a new agar plate containing 20 μg/ml erythromycin. Cells were allowed to grow for one week until colonies appeared.

### *Halomonas* construct generation

The *Halomonas* compatible construct required a substitution of the *E. coli* T7 promoter for an IPTG-inducible P_T7_-like-promoter cassette (MmP1-lacO-RiboJ-RBS; **Figure S5a-b**).^26, 46^ The gene encoding non-tagged wild-type and G462V variant CvFAP was cloned into modified pSEVA441^24^ (pHal2-CvFAP_WT_ and pHal2-CvFAP_G462V_, respectively). This plasmid contains the PT7-like-promoter, a NcoI restriction site (underlined) and a pET21b-like Shine-Delgarno sequence (SD2) upstream of the start codon (bold; TTTGTTTAACTTTAAGAAGGAGATATA<u>CC **ATGG**</u>; Supplementary Figure S6b). Both the vector and pETM11 genes were double-digested (*Nco*I and (partial) *Xho*I), gel purified then ligated and transformed into *E. coli* strain Stellar as above. Full-length construct was selected following transformation into *E. coli* strain Stellar for plasmid recircularisation, recovery and sequencing.

The insertion of *E. coli* derived plasmids into *Halomonas* strain XV12 was performed using a modified conjugation protocol.^47^ *Halomonas* constructs were transformed into *E. coli* strain S17-1^48^, and plated onto kanamycin selective LB agar. *Halomonas* strain XV12 was plated onto YTN6 agar (10 g/L tryptone, 5 g/L yeast extract; 60 g/L NaCl and 15 g/L agar), and both cultures were incubated overnight at 37 °C. Colonies of both *E. coli* S17-1 (plasmid donor) and *Halomonas* strain XV12 (recipient) were mixed together on YTN3 agar (10 g/L tryptone, 5 g/L yeast extract; 30 g/L NaCl and 15 g/L agar) and incubated overnight at 37 °C. Individual colonies were re-plated onto kanamycin-containing YTN6 agar, which is selective for *Halomonas* growth only.

### Protein expression and lysate production

Wild type CvFAP-pETM11 homologues in *E. coli* st. BL21(DE3) were cultured in LB Broth Miller (500 mL; Formedium) containing 30 μg/mL kanamycin at 37 °C with 180 rpm shaking until OD_600nm_ = 0.2. The temperature was maintained at 25 °C until OD_600nm_ = 0.6. Recombinant protein production was induced with 50 μM IPTG, and maintained at 17 °C overnight. Cells were harvested by centrifugation (8950 × *g*, 4° C, 10 min), and analysed for protein content using 12% SDS-PAGE gels (Mini-PROTEAN TGX Stain-Free Precast Gels, Bio-Rad). Protein gels were imaged using a BioRad Gel Doc EZ Imager and relative protein band intensity was determined using the BioRad ImageLab software.

Cell pellets were resuspended in lysis buffer (1.2-1.7 mL/g pellet; 50 mM Tris pH 8 containing 300 mM NaCl, 10 mM imidazole, 10% glycerol, 0.25 μg/mL lysozyme, 10 μg/mL DNase I and 1 × protease inhibitors) and sonicated for 20 minutes (20 s on, 60 s off; 30% amplitude). Cell-free lysate was prepared by centrifugation at 48000 × *g* for 30 minutes at 4 °C. Lysate samples were analysed for recombinant protein expression by SDS PAGE (12% Mini-PROTEAN-TGX stain-free gel; Bio-Rad), using Precision Plus unstained protein ladder (Bio-Rad) at 300 V for 20 minutes. Protein content was visualised using an EZ Gel Doc (Bio-Rad).

### Hydrocarbon production

*In vitro* propane production reactions (200 μL) used in the initial screen of FAP homologues and variant enzyme screens were composed of FAP-containing cell-free lysate (180 μL) and butyric acid (0.36 to 4.5 mM) in sealed glass GC vials. The reactions were incubated at 30 °C for 24 h at 180 rpm under illumination (blue LED; 455 nm). Headspace gas was analysed for propane content using a Micro GC.

*In vivo* propane production of pETM11- and pET21b-containing CvFAP_WT_ and variants in *E. coli* was performed by the following general protocol: Cultures (20-100 mL) in LB medium containing kanamycin (50 μg/mL; pETM11) or ampicillin (100 μg/mL) were incubated for 4-6 h (OD_600_ ~ 1) at 37 °C and 180 rpm, followed by induction with IPTG (100 μM) and butyric acid supplementation (1-1000 mM; pH 6.8). Triplicate aliquots (1-5 mL) each of 3 biological replicate cultures were sealed into glass vials (4-20 mL) and incubated at 30 °C for 16-18 h at 200 rpm, illuminated continuously with an LED (white or blue (455 nm or 470 nm)). Headspace gas was analysed for propane content using a Micro GC. Comparative *in vivo* studies with 10 mM butyric, isobutyric, valeric, 2-methylbutyric and isovaleric acids were performed as above, with culture induction at OD_600_ of 0.6-0.8. Data is expressed as mg hydrocarbon production per litre fermenting culture.

Photosynthetic *in vivo* butyrate and propane production studies in *Synechocystis* were performed in BG11 medium^8^ using a modified protocol as follows: Initial cultures in BG11 medium were incubated at 30 °C under 30 μE white LED until OD 720 nm reached 1.0 (~4 days). Replicate culture aliquots (2 mL) were harvested by centrifugation and re-suspended in 1 mL BG11 medium supplemented with sodium bicarbonate (150 mM), cobalt (II) nitrate hexahydrate (100 μM; for Pcoa cultures only), 50 μg/ml kanamycin, 20 μg/ml erythromycin at 30 °C +/- butyric acid (10 mM). Cultures were sealed within 4 mL gas tight vials and incubated at 30 °C for 24-48 h under blue light (average 63 μE). Headspace gas was analysed for propane content using a Micro GC, and cell-free culture supernatant samples (10 μL) were analysed for butyric acid content by HPLC using an Agilent Hi-Plex H column.

Propane production in *Halomonas* strains was performed by a modification of the *E. coli* general protocol as follows: Cultures were grown in phosphate buffered YTN6 medium (50 mM K_2_HPO_4_ pH 6.6) containing spectinomycin (pHal2-CvFAPG462V; 50 μg/mL) for 5 h at 37 °C and 180 rpm. Recombinant protein expression was induced with IPTG (0.1 mM) at a higher cell density than *E. coli* cultures (OD ~ 1.6). The remainder of the *in vivo* propane production process was performed as above, with butyric acid concentrations of 10-25 mM. The effect of cell permeabilisation was investigated by supplementing cultures with Triton X-100 (2%) and/or sucrose (1%). Butyrate transporter stimulation studies were performed in the presence of methyl and ethyl acetoacetate (0.1-30 mM). The effect of light saturation on propane production was performed by varying the distance between the cultures and the light source.

### *Halomonas* cultivation

The photobioreactor was set up in batch mode with high salt glycerol medium at pH 6.8 (5 g/L yeast extract, 1 g/L glycerol, 60 g/L NaCl, 50 μg/mL spectinomycin and 0.5 mL/L antifoam; 400 mL), pre-equilibrated at 30 °C with 60-100% stirring. An overnight starter culture (10 mL) of pHal2-CvFAP_G462V_ was added and the culture was maintained at 30 °C with an airflow rate of 1.21 L/min, automated pH maintenance, culture optical density monitoring and ambient room lighting until mid-log phase (4-5 hours). Recombinant protein expression was induced with IPTG (0.1 mM), followed by the addition of sodium butyrate (60-80 mM pH ~6.8) and blue light exposure (1625 μE), and maintained for ~48 h. During continuous flow mode, maintenance of OD_680nm_ of 1.0 was achieved by automated additions of culture medium as above. Propane production was monitored at 15 min intervals by automated headspace sampling using a Micro GC, while aqueous butyrate and glycerol depletion were detected by HPLC.

### *Synechocystis* cultivation

The photobioreactor (400 mL) was set up in batch mode with starter culture diluted 3:1 in fresh BG11+ medium (BG11 pH 8.0^37, 38^ containing TES buffer and 1 g/L sodium thiosulphate) in the presence of 150 mM NaHCO_3_. Both pH control and CO_2_ supply were maintained using 1M NaHCO3 in 2 × BG11^+^. The culture was maintained at 30 °C with maximal stirring with an airflow rate of 1.21 L/min, illumination of white LED (30 μE), automated pH maintenance (1M acetic acid in 2 × BG11^+^) and optical density monitoring (680 nm and 720 nm). After reaching an optical density of ~0.5 (720 nm), cobalt (II) nitrate hexahydrate (100 μM) was added as required, the warm white illumination was increased to 60 μE the integral actinic blue LED light panel was activated to provide 500-750 μE blue light (460 – 480 nm). The culture was maintained at 30 °C for 18-48 hours, fed and not fed respectively, with manual headspace sampling or monitoring by Micro GC to quantify propane and manual HPLC sampling from the culture to quantify butyrate.

### Analytical techniques

Propane levels were determined by manual headspace injection using an Agilent 490 Micro GC, containing an Al_2_O_3_/KCl column and a thermal conductivity detector (TCD). Headspace samples were manually introduced through a heated injector (110 °C), with an injection time of 100 ms using helium as the carrier gas (10.2 psi). During the continuous monitoring mode, fermenter exhaust gases were constantly flowing through the Micro GC cell, with periodic 100 ms sampling. Compounds were separated isothermally (100 °C) over 120 s under static pressure conditions, with a sampling frequency of 100 Hz. Propane concentrations were calculated by comparing the peak areas to a standard curve generated under the same conditions.

Aqueous culture metabolites (glycerol and butyric acid) were analysed by HPLC using an Agilent 1260 Infinity HPLC with a 1260 ALS autosampler, TCC SL column heater and a 1260 refractive index detector (RID). Cell-free culture supernatant samples (10 μL injection) were analysed isocratically on an Agilent Hi-Plex H column (300 × 7.7 mm; 5 mM H_2_SO_4_) at 60 °C with a flow rate of 0.7 mL/min for 40 minutes. Analyte concentrations were calculated by comparing the peak areas to a standard curve generated under the same HPLC conditions.

## Acknowledgements

The work was supported by C3 Biotechnologies Ltd, the UK Engineering and Physical Sciences Research Council (EP/S01778X/1; EP/J020192/1), the European Union’s Horizon 2020 research and innovation programme project PHOTOFUEL under grant agreement No 640720 and the Biotechnology and Biological Sciences Research Council (BB/M017702/1; BB/L010798/1). MA was funded by a PhD scholarship from the Newton-Mosharafa fund. EW was funded by a BBSRC Case PhD studentship. ISY was funded by a PhD scholarship from Indonesia Endowment Fund for Education (LPDP). This is a contribution from the EPSRC/BBSRC Future Biomanufacturing Research Hub and the BBSRC/EPSRC Synthetic Biology Research Centre SYNBIOCHEM.

## Author contributions

R.H. and J.M.X.H. performed the majority of the cloning and small-scale *in vivo* reactions in *E. coli* and *Halomonas*, respectively. H.S.T. performed the initial cloning and characterisation CvFAP work. E.Z.W. generated and characterised variant forms of CvFAP in *E. coli.* M.A. performed the studies to detect butane and isobutane production by FAP. I.S.Y. generated the cyanobacterial clones and M.F performed the characterisation studies. L.O.J performed molecular docking studies; S.T. assisted with the *Halomonas* fermentation studies. G.G-Q.C and P.R.J. advised on the development of *Halomonas* and *Synechocystis*, respectively, as microbial chassis. S.J.O.H. developed the LED light array and M.H.S. provided technical input into propane capture and analytics. H.S.T. and N.S.S. coordinated the project. N.S.S. directed the project and secured funding.

## Competing interests

A patent application (PCT/EP2019/060013) entitled ‘Hydrocarbon production’ is pending in relation to the production of hydrocarbon gases in engineered microbial strains. P.R.J. is a board member, and M.S. and N.S.S. are founding directors of C3 Biotechnologies Ltd.

## Materials and corresponding author

Correspondence to N. S. Scrutton

