## Supplementary Information for "Distributed Biomanufacturing of Liquefied Petroleum Gas"

[1]EPSRC/BBSRC Future Biomanufacturing Research Hub, BBSRC/EPSRC Synthetic Biology Research Centre SYNBIOCHEM Manchester Institute of Biotechnology and School of Chemistry, The University of Manchester, Manchester, M1 7DN, UK. [2]C3 Biotechnologies Ltd, The Railway Goods Yard, Middleton-in-Lonsdale, Lancashire, LA6 2NF, UK. [3]Department of Life Sciences, Imperial College London, Sir Alexander Fleming Building, London SW7 2AZ, UK. [4]School of Life Sciences, Tsinghua University, 100084 Beijing, China.

**Contents** **Page**

Additional Results and Discussion ........ 2
Supporting Figures ........ 3
Fig. S1 Comparative spectra of light sources ........ 3
Fig. S2 Custom built LED array ........ 4
Fig. S3 Expression of CvFAP variants ........ 5
Fig. S4 Effect of butyric acid concentration on propane production ........ 6
Fig. S5 *Halomonas* construct organisation ........ 7
Fig. S6 Fermentation of *Halomonas* expressing pHal2-FAP$_{G462V}$ ........ 8
Fig. S7 Fermentation of *Synechcocystis* Δ*aas* strain expressing pIY918........ 9
Supporting Tables
Table S1 Expression of FAP homologues ........ 10
Table S2 Propane production by CvFAP$_{G462V}$ variants in *E. coli*........ 11
Table S3 Molecular docking simulations ........ 12
Table S4 Propane production by CvFAP$_{G462V}$-pETM11 in *E. coli*........ 13
Table S5 Propane production by CvFAP$_{G462V}$-pET21b in *E. coli*........ 14
Table S6 Substrate specificity of CvFAP$_{G462V}$ in *E. coli*........ 14
Table S7 Effect of butyric/valeric acid blends on hydrocarbon production ........ 15
Table S8 Propane production by CvFAP$_{G462V}$ in *Halomonas*........ 16
Table S9 Oligonucleotides and other DNA sequences ........ 17
Table S10 Prefix and suffix used for DNA assembly ........ 18
Table S11 Prefix and suffix linkers ........ 19
Table S12 Synthesised protein mature amino acid sequences ........ 20
References ........ 22

### Additional Results and Discussion

#### Propane production *in vivo*

Average culture light exposure is determined by factors such as light intensity (photon flux density, PFD, average distance from the light source, culture density or opaqueness, agitation rate and the shape of the reactor vessel (cylindrical or flat bed). Initial studies *in vivo* showed that light access was a significant limiting factor, as replicate cultures with a distance deviation from the LED of even a few millimetres showed significant differences in propane yields, resulting in high calculation errors. This was confirmed by measuring the light intensity differences around each LED where the cultures were positioned. For the 470 nm LED the highest light intensity was found to be within a narrow area (9 $cm^2$) directly below the light source (615 μmol photons/s/$m^2$). However, as the cultures typically occupied a much larger area (360 $cm^2$), the average PFD was found to be considerably lower, with unacceptably high variation.

To standardise culture light exposure, we assembled a custom-built LED array light source composed of 480 individual blue LEDs, giving an area of 396 $cm^2$ of relatively consistent light intensity and a fixed average culture-to-LED distance (8 cm; **Figure S3**). The average PFD (78 $\pm$ 10 μmol photons/s/$m^2$) was similar to the average for the 470 nm LED, but importantly it showed a higher consistency of light over a wider area, and its maximal wavelength was close to the absorbance maximum of $CvFAP_{WT}$. This new light source gave greater reproducibility between replicate samples, allowing comparative studies to be performed.

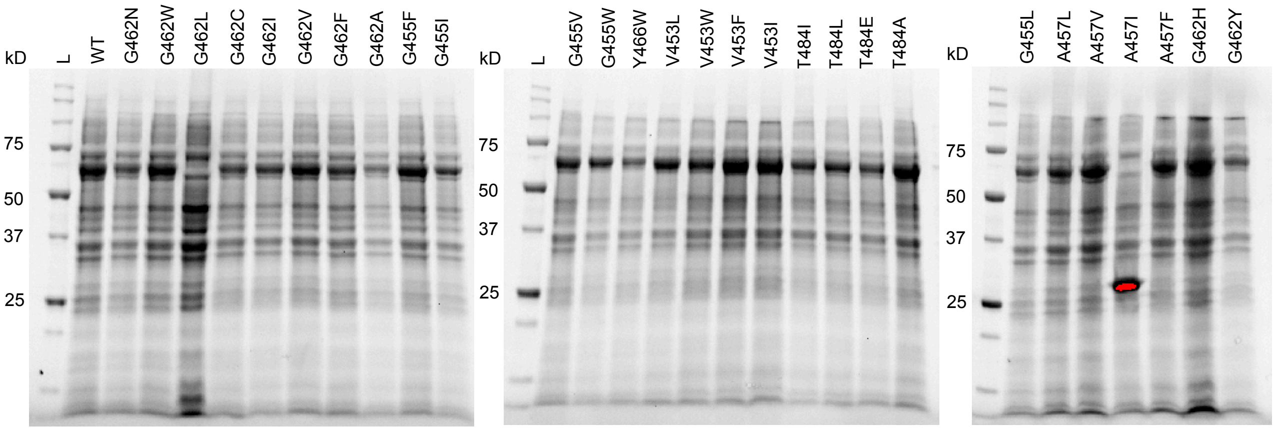


| Variant | Concentration relative to WT |
|---|---|
| WT | 1.00 |
| G462N | 0.64 |
| G462W | 1.08 |
| G462L | *No visible band* |
| G462C | 0.67 |
| G462I | 0.73 |
| G462V | 1.05 |
| G462F | 0.75 |
| G462A | 0.42 |
| G455F | 1.11 |
| G455I | 0.56 |

| Variant | Concentration relative to WT |
|---|---|
| G455V | 0.69 |
| G455W | 0.64 |
| Y466W | 0.39 |
| V453L | 0.96 |
| V453W | 0.68 |
| V453F | 1.01 |
| V453I | 0.96 |
| T484I | 0.58 |
| T484L | 0.66 |
| T484E | 0.55 |
| T484A | 1.02 |

| Variant | Concentration relative to WT |
|---|---|
| G455L | 0.39 |
| A457L | 0.65 |
| A457V | 0.90 |
| A457I | *No visible band* |
| A457F | 0.80 |
| G462H | 0.95 |
| G462Y | 0.47 |

**Figure S1. Expression of various CvFAP alleles in *E. coli*.** Protein from equal quantities of soluble cell lysate was resolved by SDS PAGE, visualized with a BioRad Gel Doc™ EZ Imager, and relative quantities of bands corresponding to CvFAP determined using instrument software. L, molecular mass ladder; WT, $CvFAP_{WT}$.

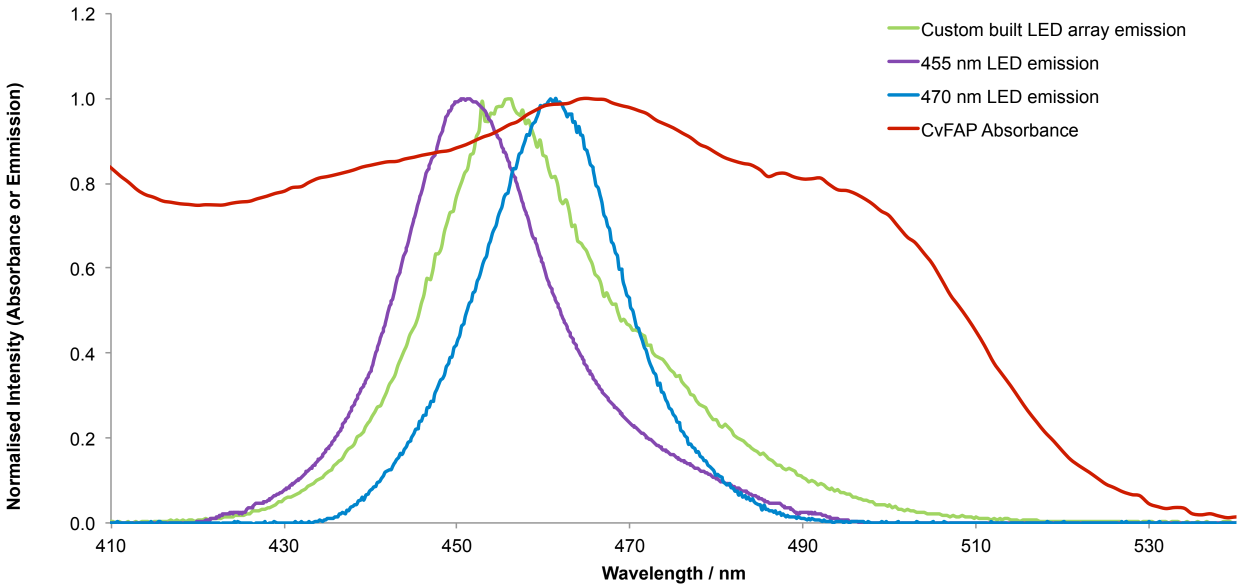


**Figure S2. Comparison of emission spectra of the three light sources with CvFAP absorbance.**

a

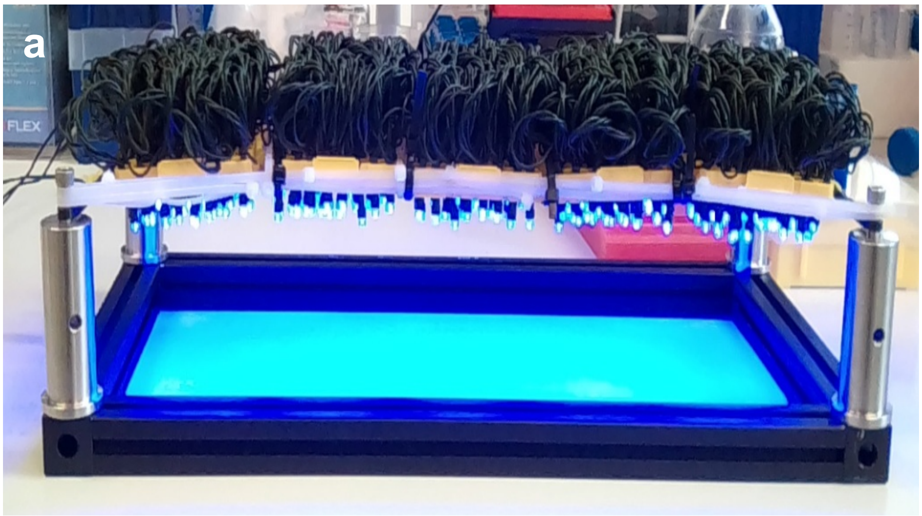

b

| | | | | | | | | | | | | |
|---|---|---|---|---|---|---|---|---|---|---|---|---|
| 31 | 37 | 40 | 39 | 38 | 41 | 43 | 44 | 40 | 39 | 40 | 39 | 26 |
| 48 | 66 | 70 | 69 | 68 | 70 | 70 | 70 | 69 | 71 | 67 | 66 | 57 |
| 62 | 72 | 84 | 87 | 89 | 87 | 87 | 84 | 93 | 89 | 90 | 79 | 63 |
| 66 | 77 | 85 | 91 | 84 | 86 | 87 | 88 | 91 | 90 | 93 | 91 | 71 |
| 50 | 61 | 66 | 68 | 68 | 69 | 67 | 70 | 75 | 74 | 77 | 61 | 57 |
| 22 | 44 | 46 | 43 | 45 | 46 | 44 | 43 | 45 | 48 | 49 | 48 | 42 |

| | | | | | | | |
|---|---|---|---|---|---|---|---|
| 0 | 20 | 40 | 60 | 80 | 100 | 200 | 300 |

**Figure S3. Custom built blue light LED array.** It is composed of 480 individual blue LEDs, giving an area of 396 $cm^2$ of relatively consistent light intensity and a fixed average culture-to-LED distance of 8 cm. Light intensity was measured with a Li-Cor light meter with a Quantum sensor, with background light value (room lights) subtracted. Values are in μmol photons/s/$m^2$ or μE. Each square is approx. 3 x 3 cm.

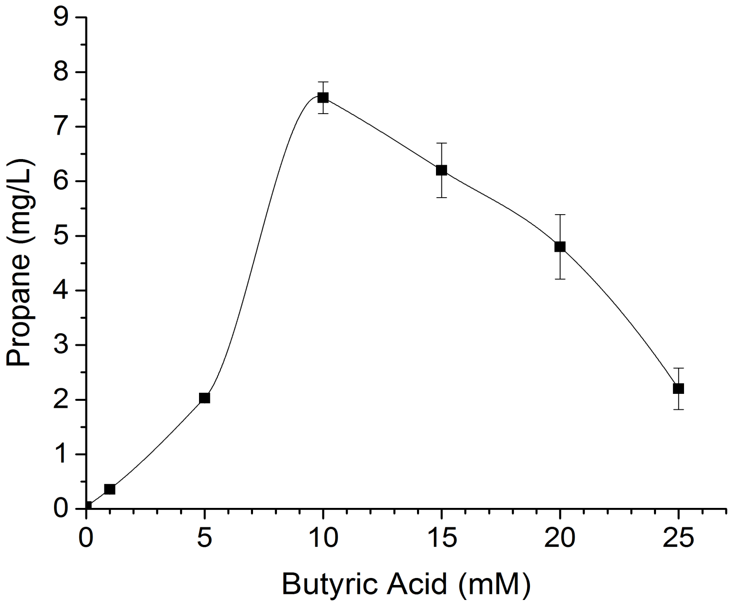


**Figure S4. Propane production by *E. coli* in increasing butyric acid concentration.** Cultures of *E. coli* BL21(DE3) containing pETm11-CvFAP$_{G462V}$ in LB broth with kanamycin (50 µg/mL) where inoculated at 1% volume from overnight starter cultures and grown for a further 6 h at 37 ºC at 180 rpm. CvFAP expression was induced with IPTG (0.1 mM), butyric acid was then added. Triplicate aliquots (1 mL each) were sealed in 5 mL glass vials and incubated at 30 ºC for 16-18 h at 200 rpm, illuminated continuously under a blue LED panel. Headspace gas was analysed for propane content using a Micro GC (100 ms injection) with an $Al_2O_3$/KCl column.

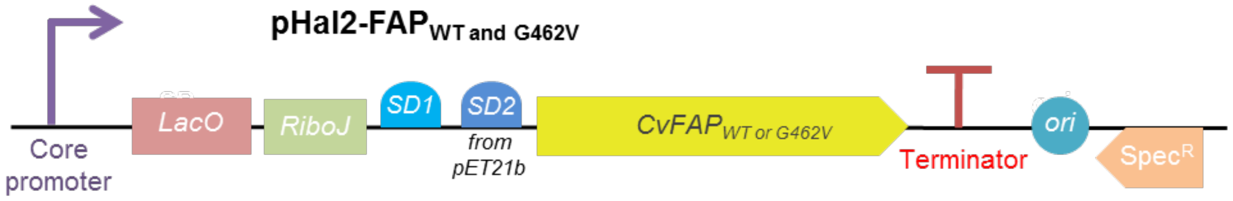


**Figure S5. Plasmid pHal2-CvFAP for expression of CvFAP in *Halomonas*.** *LacO*, *lac* operator; RiboJ$_{tr}$, truncated RiboJ (hammerhead ribozyme from the tobacco ringspot virus satellite RNA[1]); SD1 and 2, Shine-Dalgarno sequences; *ori*, origins of replication and conjugative transfer; Spec$^{R}$, spectinomycin resistance gene.

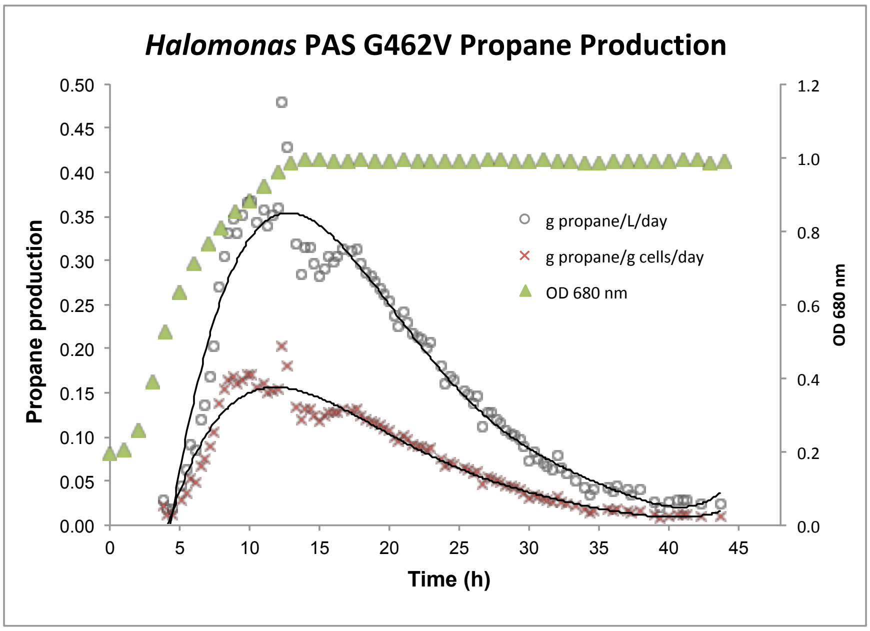


**Figure S6. Propane generation by *Halomonas* expressing pHal2-FAP$_{G462V}$ in a flat-bed photobioreactor.** Cultures were grown in high-salt glycerol medium at pH 6.8 (5 g/L yeast extract, 1 g/L glycerol, 60 g/L NaCl, 50 µg/mL spectinomycin and 0.2 mL/L antifoam; 400 mL) at 30 °C with maximal stirring and 1 L/min aeration. For crude medium, seawater with supplemental NaCl and biodiesel waste glycerol were used in place of laboratory grade reagents. CvFAP$_{G462V}$ expression was induced with IPTG (0.1 mM) at mid-log phase (indicated by an asterisk), followed by the addition of sodium butyrate (60 mM pH ~6.8) and blue light exposure (1625 µmol/s/m$^2$ photons) for up to 48 h. Culture growth was maintained at OD 680 of 1.0 by automated feed addition. Propane production was monitored every 20 minutes by automated headspace sampling using a Micro GC.

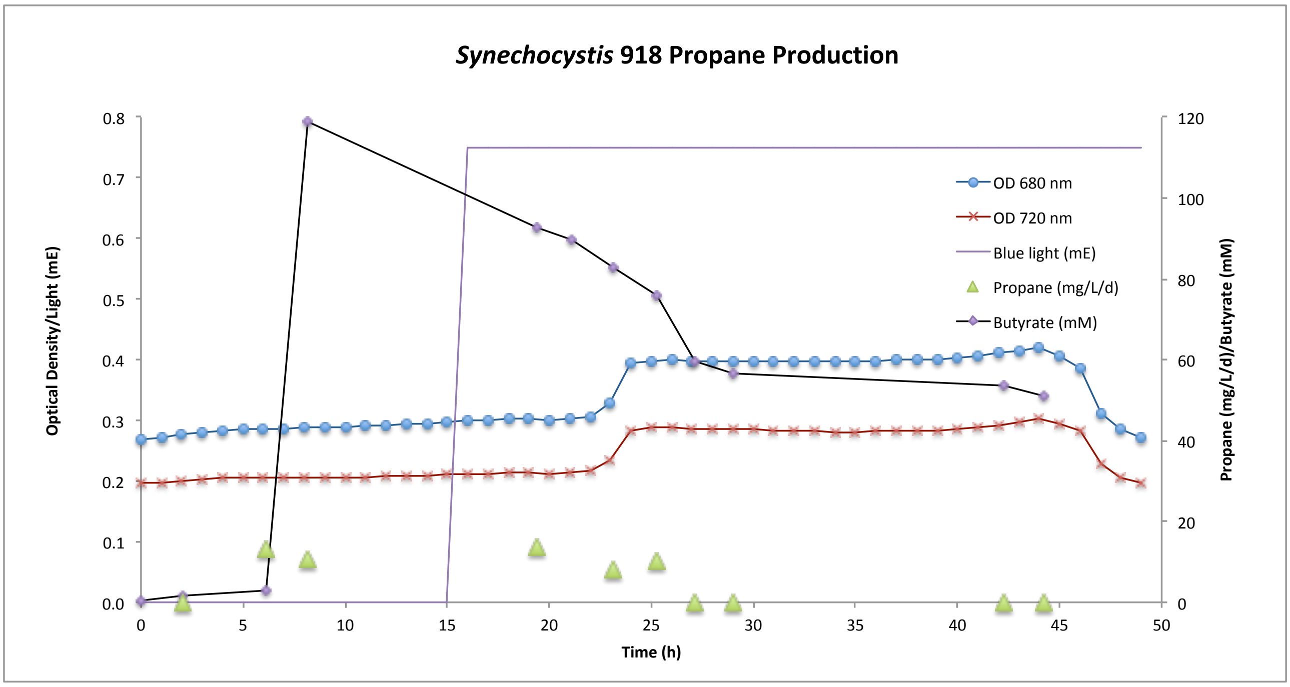


**Figure S7.** Propane generation by *Synechcocystis* Δ*aas* strain (918) with pIY918 in a flat-bed photobioreactor. The photobioreactor (400 mL) was set up in batch mode with starter culture diluted 3:1 in fresh BG11$^+$ medium (BG11 pH 8.0[2,3] containing TES buffer and 1 g/L sodium thiosulphate) in the presence of 150 mM $NaHCO_3$. Both pH control and $CO_2$ supply were maintained using 1M $NaHCO_3$ in 2 x BG11$^+$. The culture was maintained at 30 °C with maximal stirring with an airflow rate of 1.21 L/min, illumination of warm white light (30 µE), automated pH maintenance (1M acetic acid in 2 x BG11$^+$) and optical density monitoring (680 nm and 720 nm). After reaching an optical density of ~0.5 (720 nm), cobalt (II) nitrate hexahydrate (150 µM) was added as required, the warm white illumination was increased to 60 µE the integral actinic blue LED light panel was activated to provide 500-750 µE blue light (460 – 480 nm). The culture was maintained at 30 °C for 18-48 hours, fed and not fed respectively, with manual headspace sampling or monitoring by Micro GC to quantify propane and manual HPLC sampling from the culture to quantify butyrate.

### Tables

**Table S1. Expression and activity of putative FAP homologues in *E. coli*.**

| Homologue | Soluble expression | *In vitro* propane production |
|---|---|---|
| CvFAP | Very high | Yes |
| CrFAP | High | Yes |
| GpFAP | High | Yes |
| CcFAP | Low | No |
| ChFAP | Very low | No |
| CsFAP | Low | Yes |
| PtFAP | Very low | No |
| CmFAP | Low | No |

Cultures were grown in LB medium containing kanamycin (30 µg/mL) at 37 °C at 200 rpm until 0.2 $OD_{600nm}$. Recombinant protein expression was induced with IPTG (0.5 mM) and the cultures were incubated for 17 h at 17-25 °C. 180 µL of cell-free lysate was then mixed with butyric acid (400 µM in total volume 200 µL) in sealed glass GC vials. The reactions were incubated at 30 °C for 24 h at 180 rpm under a blue light LED array. Headspace gas was analysed for propane content using a Micro GC with an $Al_2O_3$/KCl column.

**Table S2.** Propane production by $CvFAP_{WT}$ and variants in *E. coli*.

| Variant | Propane production (mg/L culture) | Relative activity[1] (mg/L culture, normalised) |
|---|---|---|
| WT | 0.67 ± 0.99 | 0.67 ± 0.99 |
| V453F | 0.95 ± 0.14 | 0.94 ± 0.14 |
| V453I | 1.81 ± 0.55 | 1.89 ± 0.57 |
| V453L | 0.33 ± 0.18 | 0.40 ± 0.22 |
| V453W | 1.25 ± 0.13 | 1.85 ± 0.20 |
| G455F | 0.29 ± 0.07 | 0.26 ± 0.06 |
| G455I | 0.08 ± 0.01 | 0.15 ± 0.02 |
| G455V | 0.05 ± 0.00 | 0.07 ± 0.00 |
| G455W | 0.25 ± 0.16 | 0.41 ± 0.27 |
| G455L | 0.08 ± 0.10 | 0.21 ± 0.26 |
| A457F | 0.02 ± 0.02 | 0.02 ± 0.03 |
| A457I | 0.03 ± 0.10 | - |
| A457L | 0.04 ± 0.08 | 0.05 ± 0.13 |
| A457V | 0.07 ± 0.15 | 0.07 ± 0.17 |
| G462A | 7.14 ± 1.09 | 16.87 ± 2.58 |
| G462C | 3.94 ± 2.38 | 5.91 ± 3.57 |
| G462F | 7.00 ± 0.38 | 9.34 ± 0.51 |
| G462H | 0.04 ± 0.02 | 0.04 ± 0.02 |
| G462I | 10.77 ± 1.19 | 14.77 ± 1.63 |
| G462L | 0.02 ± 0.01 | - |
| G462N | 0.53 ± 0.08 | 0.83 ± 0.12 |
| G462V | 5.07 ± 1.12 | 4.85 ± 1.07 |
| G462W | 0.61 ± 0.19 | 0.57 ± 0.18 |
| G462Y | 0.83 ± 0.46 | 1.75 ± 0.98 |
| Y466W | 0.23 ± 0.08 | 0.60 ± 0.19 |
| T484A | 0.03 ± 0.03 | 0.03 ± 0.02 |
| T484E | 0.01 ± 0.01 | 0.02 ± 0.02 |
| T484I | 0.00 ± 0.01 | 0.01 ± 0.01 |
| T484L | 0.03 ± 0.05 | 0.04 ± 0.07 |

Cultures (20 mL) were grown in LB medium containing kanamycin (30 μg/mL) at 37 ºC until ~ 0.6-0.8 $OD_{600}$. Recombinant protein expression was induced with IPTG (0.1 mM) and cultures were supplemented with 10 mM butyric acid. Triplicate aliquots (1 mL) of cultures were sealed into 5 mL glass vials and incubated at 30 ºC for 16-18 h at 200 rpm, illuminated with a blue LED array. Headspace gas was analysed for gaseous hydrocarbon content using a Micro GC. Reactions were performed in triplicates of biological replicates. Normalised data was calculated by dividing the propane yields (mg/L culture) by the relative protein concentration compared to the wild type (WT) enzyme (**Figure S1**). [1]Lysates A457I and G462L did not show visible bands on SDS PAGE, so no relative activity data could be calculated.

**Table S3. Predicted binding affinities of CvFAP variants with butyrate and palmitate.**

| Variant | -ΔG (kcal/mol) | | $K_b$ (kcal/mol) | |
|---|---|---|---|---|
| | Butyrate | Palmitate | Butyrate | Palmitate |
| WT | 1.00 | 1.00 | 1.00 | 1.00 |
| G462V | 1.05 | 0.70 | 1.40 | 0.03 |
| G462I | 1.05 | 0.73 | 1.40 | 0.03 |
| G462L | 1.05 | 1.00 | 1.40 | 1.00 |
| V453I | 1.02 | 0.97 | 1.18 | 0.72 |
| G455I | 1.05 | 1.05 | 1.40 | 1.96 |
| Y466W | 0.97 | 0.89 | 0.85 | 0.26 |
| T484I | 1.00 | 0.95 | 1.00 | 0.51 |
| A457V | 1.03 | 0.88 | 1.18 | 0.22 |

Molecular docking simulations were performed using Autodock Vina with the wild-type crystal structure of CvFAP.[4] Values of the predicted free energy of binding (-ΔG) and binding constant ($K_b$) are normalised against the values for WT.

**Table S4.** Propane production by $CvFAP_{G462V}$ in *E. coli* in presence of additives that affect cell permeability.

| Variable | Butyric acid (mM) | Additive | Propane (mg/L culture) |
|---|---|---|---|
| Cell leakiness | 1 | None | 0.7 $\pm$ 0.05 |
| | 1 | Triton X-100 | 1.1 $\pm$ 0.07 |
| | 1 | Sucrose | 0.9 $\pm$ 0.11 |
| | 1 | Triton X-100 and sucrose | 1.0 $\pm$ 0.31 |
| Butyric acid concentration | 0 | Triton X-100 | 0.1 $\pm$ 0.01 |
| | 1 | Triton X-100 | 0.5 $\pm$ 0.01 |
| | 5 | Triton X-100 | 2.7 $\pm$ 0.02 |
| | 10 | Triton X-100 | 10.0 $\pm$ 0.39 |
| | 15 | Triton X-100 | 8.3 $\pm$ 0.66 |
| | 20 | Triton X-100 | 6.4 $\pm$ 0.79 |
| | 25 | Triton X-100 | 2.9 $\pm$ 0.51 |
| Transporter stimulation | 10 | None | 12.7 $\pm$ 0.03 |
| | 10 | 0.1 mM EAA | 12.6 $\pm$ 1.04 |
| | 10 | 1 mM EAA | 19.8 $\pm$ 1.31 |
| | 10 | 10 mM EAA | 23.4 $\pm$ 1.30 |
| | 10 | 20 mM EAA | 21.0 $\pm$ 2.31 |
| | 10 | 30 mM EAA | 8.7 $\pm$ 0.22 |
| | 10 | 10 mM MAA | 35.9 $\pm$ 8.79 |

Cultures of *E. coli* BL21(DE3) containing pETm11-$CvFAP_{G462V}$ were grown as already described (Figure S4), induced with IPTG (0.1 mM) and supplemented after a further hour with butyric acid (1 to 25 mM) $\pm$ Triton X-100 (1%) and/or sucrose (1%). Triplicate 1 mL aliquots mL of cultures were sealed in 5 mL glass vials and incubated at 30 ºC for 16-18 h at 200 rpm, illuminated continuously under a blue LED array. Headspace gas was analysed for propane content using a Micro GC (100 ms injection) with an $Al_2O_3$/KCl column. Putative small chain fatty acid transporter stimulators: EAA, ethyl acetoacetate; MAA, methyl acetoacetate.

**Table S5. Effects of small-chain fatty acid transporter stimulation on propane production by CvFAP$_{G462V}$ in *E. coli***

| Construct | Butyric acid (mM) | EAA (mM) | Propane (mg/L culture) |
|---|---|---|---|
| CvFAP$_{G462V}$ | 10 | 0 | 47.9 ± 11.8 |
| | 10 | 10 | 97.1 ± 10.3 |
| CvFAP$_{G462V}$ AtoE | 10 | 0 | 65.1 ± 3.2 |
| | 10 | 10 | 93.8 ± 8.1 |

Cultures of *E. coli* BL21(DE3) with pET21b containing genes to express either N-His$_6$-CvFAP$_{G462V}$ alone, or both N-His$_6$-CvFAP$_{G462V}$ and AtoE (a small-chain fatty acid transporter protein) were grown in LB medium containing ampicillin (100 μg/mL). Cultures were inoculated at 1% volume from overnight starter cultures and grown for 6 h at 37 ºC at 180 rpm. Recombinant protein expression was induced with IPTG (0.1 mM) and additions were made of butyric acid (10 mM) ± ethyl acetoacetate (EAA, 10 mM, a putative stimulator of small-chain fatty acid uptake). Triplicate 1 mL aliquots of cultures were sealed into 5 mL glass vials and incubated at 30 ºC for 16-18 h at 200 rpm, illuminated continuously with a blue LED panel. Headspace gas was analysed for propane content using a Micro GC (100 ms injection) with an $Al_2O_3$/KCl column.

**Table S6. Production of gaseous hydrocarbons by variant CvFAP in *E. coli* in the presence of short-chain fatty acids.**

| | Substrate acid | | | | |
|---|---|---|---|---|---|
| | Butyric | Isobutyric | Valeric | 2-MB | Isovaleric |
| Variant | Propane (mg/L culture) | | Butane (mg/L culture) | | Isobutane (mg/L culture) |
| WT | 7.0 ± 0.6 | 6.1 ± 2.4 | 17.7 ± 1.9 | 7.1 ± 1.4 | 5.6 ± 0.3 |
| G462A | 17.6 ± 0.7 | 5.0 ± 1.2 | 33.5 ± 6.5 | 50.0 ± 11.4 | 30.2 ± 3.9 |
| G462I | 43.8 ± 3.1 | 36.9 ± 5.4 | 47.1 ± 7.8 | 95.4 ± 5.8 | 86.8 ± 10.8 |
| G462F | 31.2 ± 0.7 | 31.4 ± 3.3 | 27.7 ± 0.8 | 38.5 ± 10.9 | 28.6 ± 4.0 |
| G462V | 24.5 ± 5.0 | 24.3 ± 1.6 | 21.9 ± 0.7 | 12.2 ± 2.1 | 17.4 ± 2.0 |

Cultures of *E. coli* BL21(DE3) ΔyqhD ΔyjgB with pETM11-CvFAP variants in LB medium containing kanamycin (50 μg/mL) were inoculated at 1% volume from overnight starter cultures and grown further at 37 ºC to 0.6-0.8 $OD_{600}$ Recombinant protein expression was induced with IPTG (0.1 mM) and cultures were supplemented with different shart-chain fatty acids (10 mM). Triplicate 1 mL aliquots were sealed into 5 mL glass vials and incubated at 30 ºC for 16-18 h at 200 rpm, illuminated continuously under a blue LED panel. Headspace gas was analysed for propane content using a Micro GC (100 ms injection) with an $Al_2O_3$/KCl column. WT = wild type; 2-MB = 2-methylbutyric acid.

**Table S7.** Effect of butyric/valeric acid blends on gaseous hydrocarbon production by wild type CvFAP in *E. coli* BL21(DE3) ΔyqhD ΔyjgB.

| Butyric Acid (%) | Valeric acid (%) | Propane (mg/L culture) | Butane (mg/L culture) |
|---|---|---|---|
| 0 | 0 | 0.6 ± 0.06 | 0.03 ± 0.00 |
| 0 | 100 | 0.21 ± 0.01 | 17.13 ± 0.31 |
| 20 | 80 | 3.62 ± 0.13 | 13.74 ± 0.42 |
| 30 | 70 | 5.76 ± 0.04 | 13.15 ± 0.03 |
| 35 | 65 | 6.43 ± 0.25 | 11.87 ± 0.5 |
| 40 | 60 | 7.39 ± 0.22 | 10.99 ± 0.07 |
| 50 | 50 | 8.75 ± 1.04 | 8.37 ± 0.91 |
| 60 | 40 | 11.67 ± 0.66 | 7.41 ± 0.29 |
| 70 | 30 | 11.96 ± 1.32 | 5.02 ± 0.59 |
| 80 | 20 | 14.04 ± 0.22 | 3.72 ± 0.05 |
| 90 | 10 | 17.29 ± 0.53 | 1.95 ± 0.04 |
| 92 | 8 | 17.56 ± 0.31 | 1.57 ± 0.05 |
| 95 | 5 | 17.10 ± 0.24 | 0.99 ± 0.02 |
| 100 | 0 | 19.32 ± 1.55 | 0.00 ± 0.00 |

Cultures in LB were grown and induced with IPTG (0.1 mM) as already (Table S6) described, then supplemented with butyric/valeric acid mixtures (10 mM total). Triplicate 1 mL aliquots were sealed into 5 mL glass vials and incubated at 30 ºC for 16-18 h at 200 rpm, illuminated with a blue LED panel. Headspace gas was analysed for gaseous hydrocarbon content using a Micro GC.

**Table S8. Propane production by $CvFAP_{G462V}$ in *Halomonas*.**

| Variable | Butyric acid (mM) | Additive | Propane (mg/L culture) |
|---|---|---|---|
| Cell leakiness | 10 | None | 55.2 $\pm$ 13.2 |
| | 10 | Triton X-100 | 55.5 $\pm$ 3.7 |
| | 10 | Sucrose | 50.9 $\pm$ 9.1 |
| | 25 | None | 127.8 $\pm$ 5.7 |
| | 25 | Triton X-100 | 144.6 $\pm$ 6.1 |
| | 25 | Sucrose | 121.4 $\pm$ 13.6 |
| Butyric acid concentration | 0 | None | 0.9 $\pm$ 0.1 |
| | 10 | None | 54.9 $\pm$ 1.4 |
| | 20 | None | 96.7 $\pm$ 1.6 |
| | 30 | None | 117.2 $\pm$ 15.4 |
| | 40 | None | 119.7 $\pm$ 15.3 |
| | 50 | None | 138.4 $\pm$ 6.32 |
| | 60 | None | 133.7 $\pm$ 2.25 |
| | 80 | None | 157.1 $\pm$ 17.14 |
| | 100 | None | 102.2 $\pm$ 7.0 |
| Transporter stimulation | 25 | None | 45.6 $\pm$ 1.1 |
| | 25 | EAA | 51.7 $\pm$ 0.5 |
| | 25 | MAA | 42.9 $\pm$ 1.6 |

Cultures of *Halomonas* XV12 containing pHal2-$CvFAP_{G462V}$ were grown in YTN6 medium (yeast extract 5 g/L, tryptone 10 g/L, NaCl 60 g/L, pH 9.0/NaOH) containing spectinomycin (50 μg/mL) were inoculated from overnight starter cultures at 1% volume and grown at 37 ºC at 180 rpm to 1.0–1.2 $OD_{600}$. Recombinant protein expression was induced with IPTG (0.1 mM). Cultures were adjusted to pH 6.8 by combined addition of $KH_2PO_4$ (50 mM) and butyrate (1-25 mM)*, and other additions were made as indicated (Triton X-100, 1% w/v; sucrose, 1% w/v; EAA or MAA, 10 mM). Triplicate 1 mL aliquots were sealed into 5 mL glass vials and incubated at 30 ºC for 16-18 h at 180 rpm, illuminated continuously under a blue LED array. Headspace gas was analysed for propane content using a Micro GC (100 ms injection) with an $Al_2O_3$/KCl column. EAA, ethyl acetoacetate; MAA, methyl acetoacetate.

*Optimal culture pH for propane production by CvFAP in *Halomonas* was pH 6.5-7.0. It was therefore necessary to adjust the pH of the butyrate solution accordingly prior to mixing with the culture.

**Table S9.** Oligonucleotide and other DNA sequences in *E. coli* and *Halomonas*.

| Protein | DNA sequence |
|---|---|
| OXB1 promoter[1] | AAGCTGTTGTGACCGCTTGCTCTAGCCAGCTATCGAGTTGTGAACCGATCCATCTAGCA |
| | ATTGGTCTCGATCTAGCGATAGGCTTCGATCTAGCTATGTAGAAACGCCGTGTGCTCGA |
| | TCGCCTGACGCTTTTTATCGCAACTCTCTACTGTTGCTTCAACAGAACATATTGACTAT |
| | CCGGTATTACCCGGC*CATGGTATATCTCCTTCTTAAAGTTAAACAAA* |
| *Mutagenesis in E. coli* | |
| CvFAP$_{G462V}$ | 5′-GCACTGGATCCGGAT*GTT*GTTAGCACCTATGTG-3′ |
| | 5′-CACATAGGTGCTAAC*AAC*ATCCGGATCCAGTGC-3′ |
| CvFAP$_{G462I}$ | 5′-GATCCGGATATTGTTAGCACCTATG-3′ |
| | 5′-CAGTGCCATACCAGGAACAAAAC-3′ |
| CvFAP$_{G462F}$ | 5′-GATCCGGATTTTGTTAGCACC-3′ |
| | 5′-CAGTGCCATACCAGGAACAAAAC-3′ |
| CvFAP$_{G462A}$ | 5′-GCGGTTAGCACCTATGTGCGTTTTG-3′ |
| | 5′-ATCCGGATCCAGTGCCATAC-3′ |
| CvFAP$_{G462H}$ | 5′-CAGTGCCATACCAGGAACAAAACG-3′ |
| | 5′-CAGTGCCATACCAGGAACAAAAC-3′ |
| CvFAP$_{G462L}$ | 5′-GATCCGGATCACGTTAGCACCTATG-3′ |
| | 5′-GATCCGGATCTGGTTAGCACCTATG-3′ |
| CvFAP$_{G462C}$ | 5′-GATCCGGATTGTGTTAGCACCTATG-3′ |
| | 5′-GATCCGGATTGGGTTAGCACCTATG-3′ |
| CvFAP$_{G462W}$ | 5′-GATCCGGATTATGTTAGCACCTATG-3′ |
| | 5′-GATCCGGATAACGTTAGCACCTATG-3′ |
| CvFAP$_{G462Y}$ | 5′-GATCCGGATTATGTTAGCACCTATG-3′ |
| | 5′-CAGTGCCATACCAGGAACAAAAC-3′ |
| CvFAP$_{G462N}$ | 5′-GATCCGGATAACGTTAGCACCTATG-3′ |
| | 5′-CAGTGCCATACCAGGAACAAAACG-3′ |
| CvFAP$_{G455F}$ | 5′-GTTTTGTTCCTTTTTATGGCACTGGATCC-3′ |
| | 5′-GAACTTGCAGATCCGGCAG-3′ |
| CvFAP$_{G455I}$ | 5′-GTTTTGTTCCTATTATGGCACTGGATCC-3′ |
| | 5′-GAACTTGCAGATCCGGCAG-3′ |
| CvFAP$_{G455V}$ | 5′-GTTTTGTTCCTGTTATGGCACTGGATCC-3′ |
| | 5′-GAACTTGCAGATCCGGCAG-3′ |
| CvFAP$_{G455W}$ | 5′-GTTTTGTTCCTTGGATGGCACTGGATC-3′ |
| | 5′-GAACTTGCAGATCCGGCAG-3′ |
| CvFAP$_{G455L}$ | 5′-TTTTGTTCCTCTGATGGCACTGGATCC-3′ |
| | 5′-CGAACTTGCAGATCCGGC-3′ |
| CvFAP$_{Y466W}$ | 5′-GTGTTAGCACCTGGGTGCGTTTTG-3′ |
| | 5′-CATCCGGATCCAGTGCCATAC-3′ |
| CvFAP$_{V453L}$ | 5′-CAAGTTCGTTTTCTGCCTGGTATGGCAC-3′ |
| | 5′-CAGATCCGGCAGTGCCTG-3′ |
| CvFAP$_{V453W}$ | 5′-CAAGTTCGTTTTTGGCCTGGTATGGCAC-3′ |
| | 5′-CAGATCCGGCAGTGCCTG-3′ |
| CvFAP$_{V453F}$ | 5′-CAAGTTCGTTTTTTTCCTGGTATGGCAC-3′ |
| | 5′-CAGATCCGGCAGTGCCTG-3′ |
| CvFAP$_{V453I}$ | 5′-CAAGTTCGTTTTATTCCTGGTATGGCAC-3′ |
| | 5′-CAGATCCGGCAGTGCCTG-3′ |
| CvFAP$_{T484I}$ | 5′-GCCTGAAATGGCCGAGCGGTATTDHMATGCAGCTGATTGCATGT-3′ |
| | 5′-CCTGGCTCTGAAATTTGGCAAAACG-3′ |
| CvFAP$_{T484L}$ | 5′-GCCTGAAATGGCCGAGCGGTATTDHMATGCAGCTGATTGCATGT-3′ |

| | |
|---|---|
| | 5′-CCTGGCTCTGAAATTTGGCAAAACG-3′ |
| $CvFAP_{T484E}$ | 5′-GCCTGAAATGGCCGAGCGGTATTDHMATGCAGCTGATTGCATGT-3′ |
| | 5′-CCTGGCTCTGAAATTTGGCAAAACG-3′ |
| $CvFAP_{T484A}$ | 5′-GCCTGAAATGGCCGAGCGGTATTDHMATGCAGCTGATTGCATGT-3′ |
| | 5′-CCTGGCTCTGAAATTTGGCAAAACG-3′ |
| $CvFAP_{A457L}$ | 5′-GTTCGTTTTGTTCCTGGTATGNTTCTGGATCCGGATGGTGTTAGC-3′ |
| | 5′-GCTAACACCATCCGGATCCAGAANCATACCAGGAACAAAACGAAC-3′ |
| $CvFAP_{A457V}$ | 5′-GTTCGTTTTGTTCCTGGTATGNTTCTGGATCCGGATGGTGTTAGC-3′ |
| | 5′-GCTAACACCATCCGGATCCAGAANCATACCAGGAACAAAACGAAC-3′ |
| $CvFAP_{A457I}$ | 5′-GTTCGTTTTGTTCCTGGTATGNTTCTGGATCCGGATGGTGTTAGC-3′ |
| | 5′-GCTAACACCATCCGGATCCAGAANCATACCAGGAACAAAACGAAC-3′ |
| $CvFAP_{A457F}$ | 5′-GTTCGTTTTGTTCCTGGTATGNTTCTGGATCCGGATGGTGTTAGC-3′ |
| | 5′-GCTAACACCATCCGGATCCAGAANCATACCAGGAACAAAACGAAC-3′ |
| *Assembly of FAPGV-OXB1-atoE construct in pET21b* | |
| Vector opening | 5′-CGACATCACCGATGGGGAAGA-3′ |
| | 5′-CCATCGGTGATGTCGGTCCGGCGTAGAGGATCGAG-3′ |
| Insert generation | 5′-CGGTTGCTGGCGCCTATATCTAATGCGCCGCTACAGGGC-3′ |
| | 5′-GATATAGGCGCCAGCAACCG-3′ |
| *Assembly of pHal1-$FAP_{WT}$* | |
| Vector opening | 5′-TGCCACCGCTGAGCAATAAAA-3′ |
| | 5′-CATCTAGTATTTCTCCTCTTTCTCTAGTA-3′ |
| Insert generation | 5′-GAGAAATACTAGATGGCCAGCGCAGTTGAAGATATT-3′ |
| | 5′-TGCTCAGCGGTGGCATTATGCTGCAACGGTTGCCG-3′ |
| *Generation of $FAP_{G462V}$ in pET21b* | |
| Vector opening | 5′-CTGAAAGGAGGAACTATATCCGGATTG-3′ |
| | 5′-AGTTCCTCCTTTCAGCTCTACGCCGGACGCATCGT-3′ |

[1]Italics = Shine-Delgarno sequence.

**Table S10.** Prefix and suffix used for DNA assembly.

| Assembly | Prefix linker | Plasmid | Suffix linker |
|---|---|---|---|
| Plasmid: pIY918 or pJET-Ptrc-Tes4-CvFAPG462V | | | |
| | LRBS1-4P | pIY840 | LRBS2-4S |
| 1 | LRBS2-4P | pIY882 | 1S |
| | 1P | pIY345[2] | LRBS1-4S |
| Plasmid: pIY906 or pJET-Pcoa-Tes4-CvFAPG462V | | | |
| | LRBS1-4P | pIY840 | LRBS2-4S |
| 2 | LRBS2-4P | pIY882 | 1S |
| | 1P | pIY417[2] | LRBS1-4S |
| Plasmid: pIY894 or pJET-Ptrc-CvFAPG462V | | | |
| 3 | LRBS1-4P | pIY882 | 1S |
| | 1P | pIY345 | LRBS1-4S |
| Plasmid: pIY845 or pJET-Pcoa-Tes4 | | | |
| 4 | LRBS1-4P | pIY840 | 1S |
| | 1P | pIY417[2] | LRBS1-4S |

Plasmids pIY345 and pIY417 are described in Yunus, I. S. and Jones, P. R. (2018).[2]

**Table S11.** Prefix and suffix linkers

| Adapter | | Linker | | P linker |
|---|---|---|---|---|
| **Name** | **Sequence (5' to 3')** | **Name** | **Sequence (5' to 3')** | |
| **Prefix linkers** | | | | |
| 1P-A | TTTATTGAACTA | 1P-L | GGACTAGTTCAATAAATACCC TCTGACTGTCTCGGAG | 1P |
| LRBS1-4P-A | ATCACAAGGAGGTA | LRBS1-4P-L | GGACTACCTCCTTGTGATTTA CAACTGATACTTACCTGA | LRBS1-4P |
| LRBS2-4P-A | ATCACAAGGAGGTA | LRBS2-4P-L | GGACTACCTCCTTGTGATTTT CTGCTACCCTTATCTCAG | LRBS2-4P |
| **Suffix linkers** | | | | **S linker** |
| 1S-A | TGTCGTAAGTAA | 1S-L | CTCGTTACTTACGACACTCCG AGACAGTCAGAGGGTA | 1S |
| LRBS1-4S-A | GACGGTGTTCAA | LRBS1-4S-L | CTCGTTGAACACCGTCTCAGG TAAGTATCAGTTGTAA | LRBS1-4S |
| LRBS2-4S-A | CCAATAGTAACA | LRBS2-4S-L | CTCGTGTTACTATTGGCTGAG ATAAGGGTAGCAGAAA | LRBS2-4S |

P linker = mixed prefix linker; S linker = mixed suffix linker.

**Table S12.** Putative mature amino acid sequences of the synthesised proteins.

| Protein | Amino acid sequence |
|---|---|
| CvFAP | MASAVEDIRKVLSDSSSPVAGQKYDYILVGGGTAACVLANRLSADGSKRVLVLEAGPDNTSRDV<br>KIPAAITRLFRSPLDWNLFSELQEQLAERQIYMARGRLLGGSSATNATLYHRGAAGDYDAWGVE<br>GWSSEDVLSWFVQAETNADFGPGAYHGSGGPMRVENPRYTNKQLHTAFFKAAEEVGLTPNSDFN<br>DWSHDHAGYGTFQVMQDKGTRADMYRQYLKPVLGRRNLQVLTGAAVTKVNIDQAAGKAQALGVE<br>FSTDGPTGERLSAELAPGGEVIMCAGAVHTPFLLKHSGVGPSAELKEFGIPVVSNLAGVGQNLQ<br>DQPACLTAAPVKEKYDGIAISDHIYNEKGQIRKRAIASYLLGGRGGLTSTGCDRGAFVRTAGQA<br>LPDLQVRFVPGMALDPD**G**VSTYVRFAKFQSQGLKWPSGITMQLIACRPQSTGSVGLKSADPFAP<br>PKLSPGYLTDKDGADLATLRKGIHWARDVARSSALSEYLDGELFPGSGVVSDDQIDEYIRRSIH<br>SSNAITGTCKMGNAGDSSSVVDNQLRVHGVEGLRVVDASVVPKIPGGQTGAPVVMIAERAAALL<br>TGKATIGASAAAPATVAA |
| CcFAP | MAGTVASTFRRTVPSSEAATTYDYIIVGGGAAGCVLANRLTEDPSTRVLLLEAGKPDDSFYLHV<br>PLGFPYLLGSPNDWAFVTEPEPNLANRRLYFPRGKVLGGSHAISVMLYHRGHPADYTAWAESAP<br>GWAPQDVLPYFLKSESQQSAVPNQDAHGYEGPLAVSDLARLNPMSKAFIKAAHNAAGLNHNPDF<br>NDWATGQDGVGPFQVTQRDGSRESPATSYLRAAKGRRNLTVMTGAVVERILFENPAGSSTPVAT<br>AVSFIDSKGTRVRMSASREILLCGGVYATPQLLMLSGVGPAEHLRSHGIEIVADVPAVGQNLQD<br>HAAAMVSFESQNPEKDKANSSVYYTERTGKNIGTLLNYVFRGKGPLTSPMCEAGGFAKTDPSMD<br>ACDLQLRFIPFVSEPDPYHSLADFATAGSYLQNRANRPTGFTIQSVAARPKSRGHVQLRSTDVR<br>DSMSIHGNWISNDADLKTLVHGVKLCRTIGNDDSMKEFRGRELYPGGEKVSDADIEAYIRDTCH<br>TANAMVGTCRMGIGEQAAVDPALQVKGVARLRVVDSSVMPTLPGGQSGAPTMMIAEKGADLIRA<br>AARQADAATVGAAA |
| ChFAP[1] | MAMRRLVYICAVATVTAAISSRSVPTSARRLIALRGGVAAAEQLAEEPWDYIIVGGGAAGCVMA<br>ERLSAAEARVLVLEAGTDASRDLRIRVPAGLIKVFKSERDWDFTTEAGQGTSGRGIYLCRGKAL<br>GGSSCTNVMLYNRGSPADYNSWVAAGAEGWGPDSVLHYYRKSENYVGGASQYHGVDGPLSVSDV<br>PYENELSTAFLRAAGELGYRRVHDFNDWSAPQEGFGRYKVTQRNGERCSAANAYLEGTEGRSNL<br>CVRTGVHATRVTLEGSGDDLCAAGVEYIGADGKPSRAQLAQGGEVLLSAGAVQSPQLLMLSGIG<br>PRAHLEEVGIEVRKELDNVGVGLADHPAVVVSCGSKKKVSVTDEIRLWGGSKTNPMALLRWLLW<br>RRGPLTSVACEFGGFFKTKPDLKQADVQVRFVAARAMSPDGITTLQQLGAGAKFLSGYTTQIIA<br>CRPQSTGLVRLRSSDPLAQPMLQDVHLSDDADVATLREGIKLGRQLLAAKSFDQYRDEEVYPGV<br>AVQSDEDIDAYVRKTTHSANALVGSCRMGRVDDQAAVLDPEMRVRGVGSLRVVDASAMPHIIGG<br>QTCGPTIMMAEKAADLVLRQRAEINAYMQQAQAYLAASAGAATPALSPAQAA |
| CmFAP | MAQYDFIIVGAGAAGCVLANRLSTAQFSNGDRRYPRVLLLEAGDALAEAPYFEHIPLGFPQLIG<br>SRLDYGFFSRENPTHLGGRGAVYLPRGRGEGGSHAISVMLVHRGSRHDYETWVKDYEALGWGPD<br>DVLPYFKRLESNERTAQRGADGEAATALHGSDGPLRVSDQRSPNPLSLAFIEACLERGIRRNKD<br>FNDWDHGQEGAGLFQVTQRDGRRESPATAYLQPVRSRRNLHIETNALAEHLVWSKDGRRVEGIR<br>FIDRHGRRRAALAHCEVILAAGAINTPQLLMLSGLGPGAHLQDFGIPVVRDLPGVGQNLQDHAA<br>VMLSYYAPDPYGKDRDKKRIFYTERLGKDPLVLAEYFLLGRGPLTSPVCEAGAFVHTQAVIGEP<br>SCDLQLRFVPFFSDADPYKSLGEYRSGGHVLTNTSIRPAGFGLQAVAIRPRSRGRIELATIDPR<br>ARPIIHTGWLEDKRDLQTLLSGLKLGREILSGDSMRPYRGREAFPETLEDDLVTYIRRTCHTAN<br>AIVGTARMGTGRDAVVDPELRVHGVERLRVIDASVMPKIIGGQTGVPTMMIAERGADLVKKTWK<br>LV |
| CrFAP | MASVRAAAGPAGSEKFDYVLVGGGTASCVLANKLSADGNKKVLVLEAGPTGDAMEVAVPAGITR<br>LFAHPVMDWGMSSLTQKQLVAREIYLARGRMLGGSSGSNATLYHRGSAADYDAWGLEGWSSKDV<br>LDWFVKAECYADGPKPYHGTGGSMNTEQPRYENVLHDEFFKAAAATGLPANPDFNDWSHPQDGF<br>GEFQVSQKKGQRADTYRTYLKPAMARGNLKVVIGARATKVNIEKGSSGARTTGVEYAMQQFGDR<br>FTAELAPGGEVLMCSGAVHTPHLLMLSGVGPAATLKEHGIDVVSDLSGVGQNLQDHPAAVLAAR<br>AKPEFEKLSVTSEVYDDKCNIKLGAVAQYLFQRRGPLATTGCDHGAFVRTSSSLSQPDLQMRFV<br>PGCALDPDGVKSYIVFGELKKQGRAWPGGITLQLLAIRAKSKGSIGLKAADPFINPAININYFS |

| | |
|---|---|
| | DPADLATLVNAVKMARKIAAQEPLKKYLQEETFPGERASSDKDLEEYIRRTVHSGNALVGTAAM<br>GASPAAGAVVSSADLKVFGVEGLRVVDASVLPRIPGGQTGAATVMVAERAAALLRGQATIAPSR<br>QPVAV |
| CsFAP | MAPAADKYDFILVGGGTAGCVLANRLTADGSKKVLLLEAGGANKAREVRTPAGLPRLFKSALDW<br>NLYSSLQQAASDRSIYLARGKLLGGSSATNATLYHRGTAADYDAWGVPGWTSQDALRWFIQAEN<br>NCRGIEDGVHGTGGLMRVENPRYNNPLHEVFFQAAKQAGLPENDNFNNWGRSQAGYGEFQVTHS<br>KGERADCFRMYLEPVMGRSNLTVLTGAKTLKIETEKSGGATVSRGVTFQVNGQDGSKHSAELAA<br>GGEVVLCAGSIHSPQILQLSGIGPQAELRSKDIPVVADLPGVGQNMQDHPACLSAFYLKESAGP<br>ISVTDELLHTNGRIRARAILKYLLFKKGPLATTGCDHGAFVKTAGQSEPDLQIRFVPGLALDPD<br>GIGSYTAFGKMKDQKWPSGITFQLLGVRPKSRGSVGLRSDDPWDAPKLDIGFLTDKEGADLATL<br>RSGIKLSREIAAEPAFGAYVGNELHPGAAASSDSAIDSFIRDTVHSGNANVGTCSMGVNGNAVV<br>DPSLRVFGIRGLRVADASVIPVIPGGQTGAATVMVAERAAEILLGSNQKQPAAAVPAAQPALA |
| GpFAP | MAPVDPAEKYDYILVGGGTAGCVLANKLSADGNKKVLVLEAGPSGDSLEVAVPAGIARLFAHPV<br>MDWGMSSLTQKQLVAREIYLARGRLLGGSSGTNATLYHRGTSSDYDSWGLEGWTSKDVLDWFVK<br>AECYGDGPKPYHGNSGSMNVEQPRYQNPLHEEFFRAAAAAGIPANPDFNDWSRPQDGYGEFQVA<br>QNKGQRADTYRTYLKPALSRGNLKVVTGARTTKVHIEKGSSGPRARGVEFATQQFGDRYSAQLA<br>PGGEVLMCTGAVHTPHLLMLSGVGPAAALREHGVDVVADLAGVGANLQDHPAAVVAVRAKPEFE<br>KLSVTSEIYDEKCNIKLGAVAQYLFNRRGPLATTGCDHGAFVRTSGSHSQPDLQMRFVPGCALD<br>PDGVKSYIVFGELKKQGRAWPGGITLQLLAIRAKSKGSIGLKAADPFINPAININYFSDPADLA<br>TLKQGVRMARDIARQEPLRKYLQEETFPGERASSDSDIEEYVRRTVHSGNALVGTCAMGTSPAK<br>GAVVSSSDLKVFGVEGLRVVDASVLPQIPGGQTGAATVMVAERAAALLKGQTTMAPSRQPVAA |
| PtFAP[1] | MAYDYIICGGGLAGCVLAERLSQDESKRVLVLEAGGSDYKSLFIRIPAGVLRLFRSKYDWQHET<br>GGEKGCNGRNVFLQRGKILGGSSCTNVCLHHRGSAEDYNSWNIPGWTATDVLPFFKQSQKDETG<br>RDATFHGADGEWVMDEVRYQNPLSKLFLEVGEAAGLGTNDDFNNWSHPQDGVGRFQVSEVNGER<br>CSGATAFLSKAAKRSNVIVRTGTMVRRIDFDETKTAKGITYDLMGDDTCTVPCLKEGGEVLVTG<br>GAIASPQLLMCSGIGPGKHLRSLGIPVVHDNSAVGENLQDHPAAVVSFKTPQKGVSVTSKLRLF<br>GKTNPIPVFQWLFFKSGLLTSTGCDHGAFVRTSDSLEQPDLQIRFLAARALGPDGMTTYTKFRT<br>MKTVEDGYSFQSVACRAKSKGRIRLSSSNSHVKPMIDGGYLSNQDDLATLRAGIKLGRMLGNRP<br>EWGEYLGQEVYPGPDVQTDEEIDEYIRNSLHTANALTGTCKMGTGRGAVVGPDLRVIGVNGVRV<br>ADSSVFPCIPGGQTATPTVMIADRAAVFVR |
| atoE[2] | MIGRISRFMTRFVSRWLPDPLIFAMLLTLLTFVIALWLTPQTPISMVKMWGDGFWNLLAFGMQM<br>ALIIVTGHALASSAPVKSLLRTAASAAKTPVQGVMLVTFFGSVACVINWGFGLVVGAMFAREVA<br>RRVPGSDYPLLIACAYIGFLTWGGGFSGSMPLLAATPGNPVEHIAGLIPVGDTLFSGFNIFITV<br>ALIVVMPFITRMMMPKPSDVVSIDPKLLMEEADFQKQLPKDAPPSERLEESRILTLIIGALGIA<br>YLAMYFSEHGFNITINTVNLMFMIAGLLLHKTPMAYMRAISAAARSTAGILVQFPFYAGIQLMM<br>EHSGLGGLITEFFINVANKDTFPVMTFFSSALINFAVPSGGGHWVIQGPFVIPAAQALGADLGK<br>SVMAIAYGEQWMNMAQPFWALPALAIAGLGVRDIMGYCITALLFSGVIFVIGLTLF |

[1]No chloroplast or mitochondrial targeting sequence identified. [2]Full length genes. Red = mutation to generate a N-terminal *Nco*I restriction site; Orange bold = location of the G462V mutation.
